# Single-cell analyses of the corneal epithelium: Unique cell types and gene expression profiles

**DOI:** 10.1101/2020.08.06.240036

**Authors:** Surabhi Sonam, Sushant Bangru, Kimberly J. Perry, Auinash Kalsotra, Jonathan J. Henry

**Affiliations:** Department of Cell and Developmental Biology, University of Illinois, Urbana-Champaign, IL; Department of Biochemistry, University of Illinois, Urbana-Champaign, IL; Carl R. Woese Institute for Genomic Biology, University of Illinois, Urbana-Champaign, IL; Cancer Center@Illinois, University of Illinois, Urbana-Champaign, IL

**Keywords:** Cornea, Single-cell RNA-sequencing, *Xenopus*, Stem cells, Differentiation, Biomarkers

## Abstract

Corneal Epithelial Stem Cells (CESCs) and their proliferative progeny, the Transit Amplifying Cells (TACs), are responsible for homeostasis and maintaining corneal transparency. Owing to our limited knowledge of cell fates and gene activity within the cornea, the search for unique markers to identify and isolate these cells remains crucial for ocular surface reconstruction. We performed single-cell RNA sequencing of corneal epithelial cells from stage 49-51 *Xenopus* larvae. We identified five main clusters with distinct molecular signatures, which represent apical, basal and keratocyte cell types as well as two discrete proliferative cell types in the bi-layered epithelium. Our data reveal several novel genes expressed in corneal cells and spatiotemporal changes in gene expression during corneal differentiation. Through gene regulatory network analysis, we identified key developmental gene regulons, which guide these different cell states. Our study offers a detailed atlas of single-cell transcriptomes in the frog corneal epithelium. In future, this work will be useful to elucidate the function of novel genes in corneal homeostasis, wound healing and cornea regeneration, which includes lens regeneration in *Xenopus*.

**SUMMARY STATEMENT:** This study identifies cell types and transcriptional heterogeneity in the corneal epithelium that regulate its differentiation, and facilitates the search for corneal stem cell markers.

## INTRODUCTION

The corneal epithelium is the outermost layer of the eye, and is essential for focusing light, maintaining normal vision and protecting deeper eye tissues. Owing to its constant exposure to the external environment, the corneal epithelial tissue has a remarkable capacity for self-renewal and repair, which is attributed to the presence of a population of somatic stem cells, called Corneal Epithelial Stem Cells (CESCs), and their proliferative progeny, the Transit Amplifying Cells (TACs). Therefore, the self-renewing corneal epithelium serves as an outstanding model to study the biology of epithelial stem cells, and their contributions to a highly specialized differentiated tissue.

The cellular composition and homeostatic maintenance of the corneal epithelium have been studied extensively (Cotsarelis et al., 1989; Di Girolamo, 2011; Lavker et al., 2004; Schermer et al., 1986; Thoft and Friend, 1983). In adult vertebrates, the stratified corneal epithelium consists of an innermost layer of undifferentiated basal cells, suprabasal cells, and differentiated, superficial (apical) squamous cells. Some studies in humans and mice (Amitai-Lange et al., 2015; Davanger and Evensen, 1971; Di Girolamo, 2011; Zhao et al., 2009) provide evidence that CESCs reside in the basal epithelium (in the specialized niche called the “limbus”). Contrary to this, other studies suggest that CESCs are distributed throughout the basal corneal epithelial layer (Chang et al., 2008; Majo et al., 2008). Despite some ambiguity about their location, it is known that these cells divide either symmetrically for stem cell renewal, or asymmetrically to produce more highly proliferative TACs that migrate centripetally to populate the corneal epithelium (Kinoshita et al., 1981; Tseng, 1989). As these TACs divide and migrate superficially, they progressively become more differentiated and eventually give rise to post-mitotic, terminally differentiated cells (TDCs), which undergo senescence and are sloughed off from the surface. Despite the long known existence of CESCs and TACs among vertebrates, we are limited in our understanding of the comprehensive genetic profiles of these cells and the maturation of their differentiated progeny. This has largely affected the search for definitive molecular markers to identify, isolate and enrich corneal stem cells, which are used for clinical applications.

Several studies on transcriptional profiling of corneal epithelial cells isolated from humans and mice have relied on bulk-sampling techniques and cell enrichment using pre-defined markers (Bath et al., 2013; Ma and Lwigale, 2019; Sartaj et al., 2017; Zhou et al., 2006). As these studies were restricted to specific subpopulations or regions of the corneal epithelium, it has been difficult to compare findings across these studies and systematically analyze corneal epithelial heterogeneity. Therefore, to facilitate the search of corneal stem cell biomarkers, one needs to characterize the transcriptome of many individual corneal epithelial cells representing progressive stages of vertebrate corneal cell maturation.

The South African clawed frog, *Xenopus laevis*, has been a valuable research model, particularly for studying eye tissue development, repair and regeneration, which includes the cornea and lens (Barbosa-Sabanero et al., 2012; Henry et al., 2008; Kha et al., 2018; Tseng, 2017). The anatomy and development of the frog cornea are nearly identical to that of humans (Hu et al., 2013) making it an excellent, low-cost vertebrate model to study cornea biology and determine the dynamics of corneal stem cells. Early during development, the larval corneal epithelium consists of a single basal layer, a distinct apical/superficial layer of differentiated cells and relatively few keratocytes underly this epithelium (e.g., stages 48-51) (Nieuwkoop and Faber, 1956). As the larvae undergo metamorphosis and mature into an adult (developmental stage 66), the cornea undergoes dramatic changes—the stromal space thickens and is filled with collagen lamellae and keratocytes, new epithelial layers are added, and limbal crypt-like palisades are observed (similar to those seen in mammals) (Goldberg and Bron, 1982; Hamilton et al., 2016). The presence of basal proliferative stem cells and TACs in the frog cornea has been demonstrated in previous studies. For example, cells undergoing S phase DNA replication and mitosis (examined by EdU labeling and anti-phospho-Histone H3 (S10)) are scattered throughout the basal corneal epithelium of tadpoles (Perry et al., 2013; Thomas and Henry, 2014). Whereas in the adult frog cornea, long-term BrdU, or EdU label-retaining cells, that likely identify CESCs, become concentrated in the limbal area (Hamilton and Henry, 2016). Furthermore, molecular characterization of corneal epithelia of larval and adult frogs, reveals that putative stem cell markers reported in other vertebrates are expressed at different stages in different epithelial layers of the frog cornea (Sonam et al., 2019). Although these markers label different cellular layers and regions of the corneal epithelium during maturation, none was found to be unique to CESCs or TACs. This is in general agreement with the conclusions reached by immunolocalization studies using corneas of other vertebrates (Kammergruber et al., 2019; Morita et al., 2015; Mort et al., 2012; Nowell and Radtke, 2017; Schlotzer-Schrehardt and Kruse, 2005). However, these studies provide only a glimpse into the genes expressed by these cells. Single-cell resolution molecular data is needed to provide a comprehensive, unbiased picture of tissue heterogeneity and to better understand the cellular states and transcriptional profiles during corneal differentiation.

Single-cell RNA sequencing (scRNA-seq) has emerged as a revolutionary tool that allows simultaneous, unbiased sampling of individual cells for high-throughput gene expression profile analysis to determine the cellular, as well as molecular heterogeneity in various tissues and organs (Zheng et al., 2017). This next-generation sequencing technology has enabled the identification of cell subpopulations, analysis of rare cell types, reconstruction of cell lineage trajectories and the building of gene regulatory networks for tissues and organs including the skin, heart, kidney, uterine epithelium and the intestinal epithelium (Griffiths et al., 2018; Haber et al., 2017; Haensel et al., 2020; Joost et al., 2016; Lescroart et al., 2018; Park et al., 2018; Wu et al., 2017). scRNA-seq has been particularly helpful for gaining insights into the biology and transcriptional states of different types of stem cells, including hematopoietic progenitors, neural progenitors, embryonic stem cells, muscle stem cells and others (Dell’Orso et al., 2019; Klein et al., 2015; Kumar et al., 2017; Llorens-Bobadilla et al., 2015; Zhou et al., 2016). To date, a few studies have used scRNA-seq to gain insights into the molecular and cellular heterogeneity of eye tissues, including human and mouse retina (Macosko et al., 2015; Voigt et al., 2019) and mouse cornea (Kaplan et al., 2019).

Here, we use scRNA-seq to expand our understanding of the vertebrate corneal epithelium in an amphibian model, having undertaken a comprehensive characterization of cellular and transcriptional heterogeneity to identify novel corneal marker genes. By examining 4,810 epithelial cells of tadpole corneas, we identified five distinct cell clusters, each having discrete transcriptional signatures. We used immunofluorescence studies to spatially map the scRNA-seq molecular heterogeneities (for specific genes, such as *cldn4, krt8.2*) in the intact corneal tissue. In order to better understand the dynamics of corneal cells during maturation, we also reconstructed corneal differentiation by ordering epithelial cells along a pseudo-temporal trajectory. Furthermore, our analysis of gene regulatory networks helped uncover the dynamic activity of key regulons within the corneal epithelium, that control proliferation and differentiation within this tissue. This is the first study to uncover the transcriptional profile of epithelial cells in the larval frog cornea. The *Xenopus* corneal epithelium is characterized by robust regenerative capacity and is also the source of regenerated lenses (Freeman, 1963; Perry et al., 2013). This work sets the stage for future scRNA-seq research to examine gene expression profiles during various wound repair and regeneration stages, as well as discern the cellular source and precise role of specific genes involved in corneal cell differentiation and lens regeneration.

## RESULTS

### Single-cell RNA sequencing identifies the major cell types in the tadpole corneal epithelium

To study the cellular and transcriptional heterogeneity during the early stages of cornea development, we isolated corneal epithelial cells from frog larvae. We developed an Accutase enzyme-based digestion method for dissociating single cells, pooled from 80 larval corneal epithelia (**Fig. 1A**). Using the 10X Genomics-based scRNA-seq platform, we sequenced 5,173 cells with an average of 83,449 reads per cell and the median number of genes detected per cell was 2,559 (**Table S2**). For quality control and filtering of reads, a standard computational pipeline in Seurat was used, following which 4,810 high-quality cells were retained and used for downstream computational analysis (**Fig. 1B** and **Fig. S1**). Next, an unbiased, graph-based clustering algorithm of the Seurat software was used to group cells according to their gene expression profiles. We identified five distinct cell clusters, visualized in two-dimensional space using Uniform Manifold Approximation and Projection (UMAP) (Becht et al., 2018) (**Fig. 1C**).

**Fig. 1.**
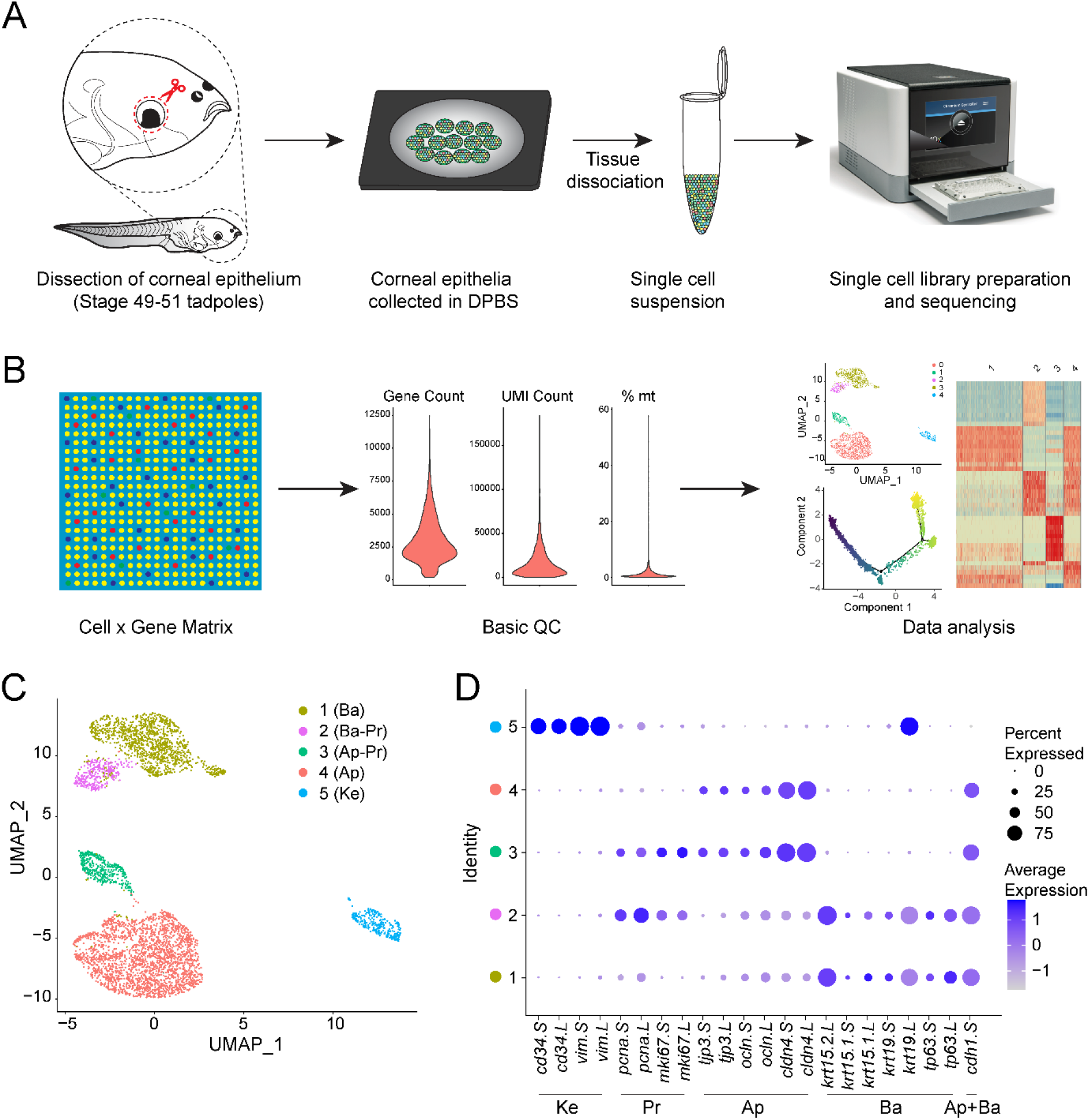
Single-cell RNA sequencing analysis of corneal cells from *Xenopus laevis* tadpoles. (**A**) Overview detailing physical workflow for isolation of corneal epithelial cells for single-cell RNA sequencing (scRNA-seq). Corneal epithelial tissues were dissected from each tadpole, collectively pooled (in DPBS) and enzymatically dissociated to isolate single corneal cells. After determining cell viability, a single-cell library was prepared from the suspension using the 10X Chromium Single Cell 3’ Reagent Kit (V3 chemistry). (**B**) Computational workflow depicting the data processing and analysis pipeline for scRNA-seq data. Cell Ranger was used to align raw reads and generate feature-barcode matrices from the scRNA-seq output. Seurat v3.1 was used to perform basic QC (see **Table S2**). (**C**) Unbiased clustering of 4,810 high-quality cells after QC cutoffs, visualized by Uniform Manifold Approximation and Projection (UMAP). Each dot represents a single cell, and cells belonging to the same cluster are similarly colored, as indicated by the key. (**D**) Dot plot showing expression of known cell-type marker genes across cells for each cluster (cell type) identified in the dataset. Dot diameter depicts the percentage of cluster cells expressing that marker and the color intensity encodes average expression of a gene among all cells within that cluster, according to the keys. Abbreviations: Ap: Apical; Ba: Basal; Ke: Keratocytes; Pr: Proliferative; Ap-Pr: Apical-Proliferative; Ba-Pr: Basal Proliferative; Ap+Ba: Apical and Basal.

To broadly attribute cell identities to these clusters, expression of known cell-type-specific marker genes reported from corneal studies in various vertebrates (Castro-Muñozledo, 2015; Chen et al., 2004; Davies et al., 2009; Yam et al., 2020; Yi et al., 2000) was compared across clusters (**Fig. 1D**). In addition, the clusters were examined for biomarkers that we previously characterized in larval *Xenopus* corneas (Sonam et al., 2019) (**Fig. 2**). Clusters 1 and 2 are comprised of basal epithelial cells. The identity of basal cells was established by the localized expression of *tp63.L*, *tp63.S*, *krt19.L*, *krt19.S*, *pax6.L*, *itgb1.L* and *itgb1.S* marker genes (**Fig. 2A-G**), which we immunolocalized to the larval corneal basal epithelium in our earlier work. Likewise, other known basal epithelial markers, including *txnip.L*, *fzd7.L*, *krt15.1.L*, *krt15.1.S*, and *krt15.2.L*, were also enriched in these two clusters, further confirming their identity (**Fig. S2A**). Cells in clusters 3 and 4 expressed markers of differentiated epithelial cells (*tjp3.L*, *tjp3.S*, *ocln.L*, *ocln.S*, *gjb1.L* and *gjb1.S*), known to be associated with apical cellular layers (**Fig. S2B**). Previously, we found beta-catenin as a marker specific for apical epithelial cells in the tadpole cornea (Sonam et al., 2019). However, in the current transcript dataset, the expression of *ctnnb.L* and *ctnnb.S* genes was detected in basal cells as well (**Fig. 2H-I**). As this observation does not fully agree with our earlier immunolocalization result, the differences could be related to those in protein and transcript expression. We also identified that marker gene *e-cadherin* (*cdh1.S*) was expressed in both basal and apical cells (Clusters 1-4; **Fig. 2J**). This follows our prior observation of E-Cadherin, as a uniform epithelial cell membrane marker in tadpole corneas (Sonam et al., 2019). Markers associated with Cluster 5 cells (*vim.L*, *vim.S*, *cd34.L*, *cd34.S* and *fn1.S*) were molecularly very distinct from the other cell clusters and we classified this cluster as corneal keratocytes (**Fig. 2K-L** and **Fig. S2C**). The presence of closely associated keratocytes underneath the larval corneal epithelium has been previously demonstrated by immunostaining (Sonam et al., 2019). We did not detect significant expression for known marker genes associated with other ocular cell types, such as *zp4*, *pou6f2*, *pitx2*—markers for corneal endothelial cells (Yoshihara et al., 2017; Yoshihara et al., 2015) and *cryaa*, *cryab*—markers for lens cells (Brahma and McDevitt, 1974; Henry et al., 2002), indicating absence of these cell types (**Fig. S3A-B**). The expression for *tcf7l2*—a marker gene that differentiates cornea versus surrounding skin epithelium (Sonam et al., 2019) was observed within very few cells, suggesting possible incorporation of some adjacent skin during corneal surgical isolation (**Fig. S3C**). On the other hand, no expression of skin melanophore-specific genes (e.g., *tyr*, *dct*, *sox10*, *tbx2*) was detected (**Fig. S4**).

**Fig. 2.**
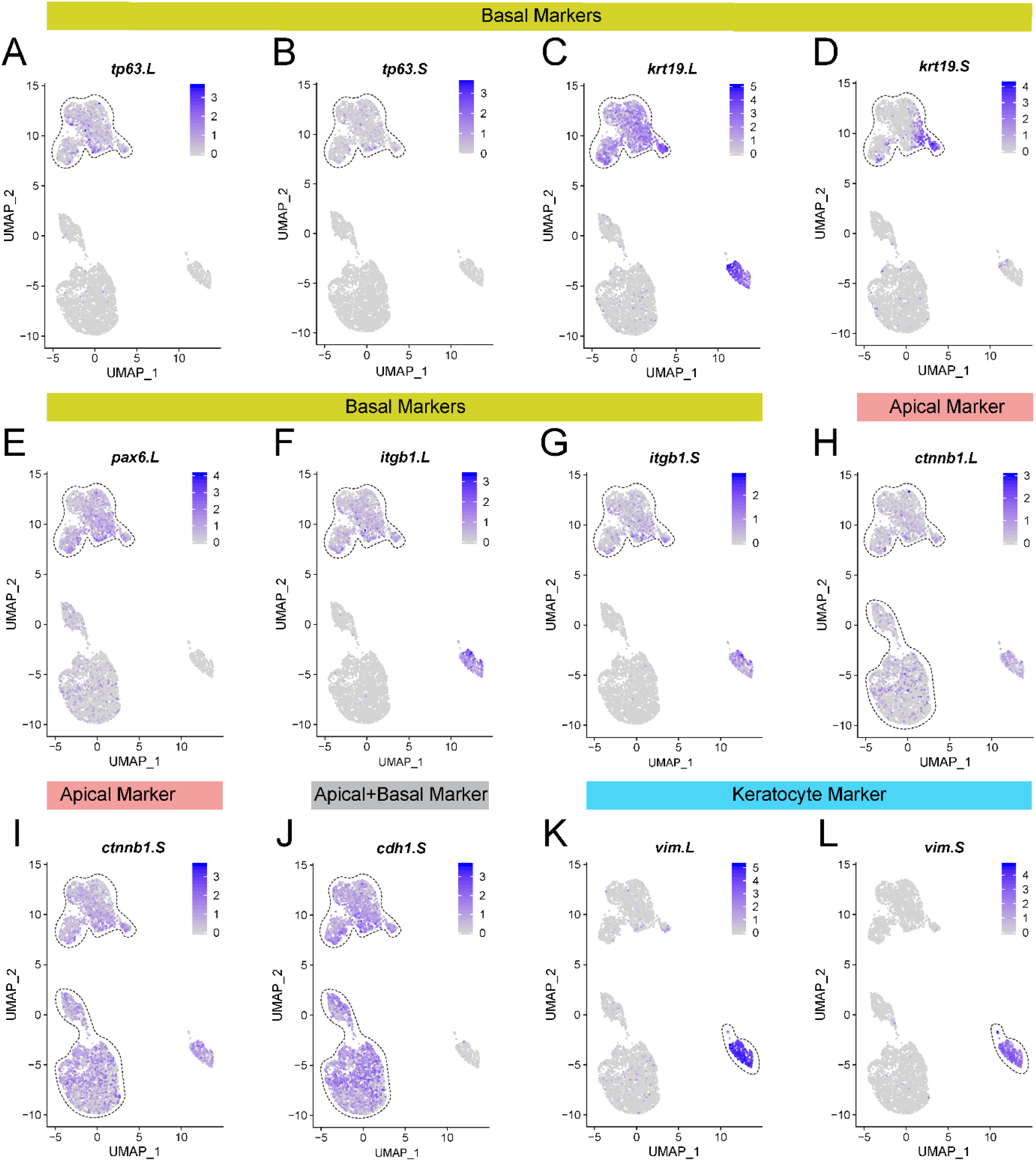
Cellular heterogeneity of larval frog corneal epithelium. Feature plots representing gene expression of previously identified (Sonam et al., 2019) *Xenopus* cell-type-specific markers that were used for assigning cluster identities. Black dashed lines highlight clusters with enhanced expression for the indicated genes. (**A-G**) Basal epithelial cells (Clusters 1 and 2) were identified based on the expression of genes *tp63.L*, *tp63.S*, *krt19.L*, *krt19.S*, *pax6.L*, *itgb1.L* and *itgb1.S*. (**H-I**) Gene *beta-catenin* (*ctnnb1.L* and *ctnnb1.S*) was used as an apical marker based on our previous immunolocalization study; however, transcripts were found to be expressed in both apical and basal cells (Clusters 1-4). (**J**) *cdh1.S* was expressed in both apical and basal cells (Clusters 1-4). (**K-L**) Corneal keratocytes (Cluster 5) were identified based on the expression of *vim.L* and *vim.S*. Color intensity correlates with the relative transcript level for the indicated gene in each cell, as indicated by the key. Additional marker gene plots are provided in **Fig. S2**.

### Proliferative cells characterize both epithelial layers of the larval cornea

A computational cell cycle analysis was performed to determine the proliferative nature of each cluster (**Fig. 3A**). We identified that clusters 2 and 3 have a proliferative characteristic, with enriched expression of proliferation marker genes including *mki67.L*, *mki67.S, pcna.L*, *pcna.L* and *top2a.L* (**Fig. 3B-F**). In contrast, clusters 1, 4 and 5 display a more quiescent nature. To verify the presence of proliferative cells in both epithelial layers, we immunostained the larval cornea using the ubiquitous cell cycle marker PCNA. Using a *Xenopus* specific antibody (Agathocleous et al., 2009), we detected PCNA labeled nuclei in both basal and apical epithelia (**Fig. 3G-H, G’-H’**). The presence of proliferative cells in both epithelial layers differs from previous *Xenopus* corneal studies (Hu et al., 2013; Perry et al., 2013), reporting the presence of dividing cells in the basal epithelium alone. The proliferative cells in the corneal epithelium are those involved in continuous growth and renewal of the tadpole cornea. To support our observation, we immunostained the corneal tissue with an anti-phospho-histone-H3S10 antibody and detected the presence of labeled nuclei in both the epithelial layers (**Fig. 3I-J. I’-J’**). As phosphorylated Histone H3 is a marker of mitotic cells (Hans and Dimitrov, 2001), the positive staining further reveals cell division activity in this bi-layered epithelium.

**Fig. 3.**
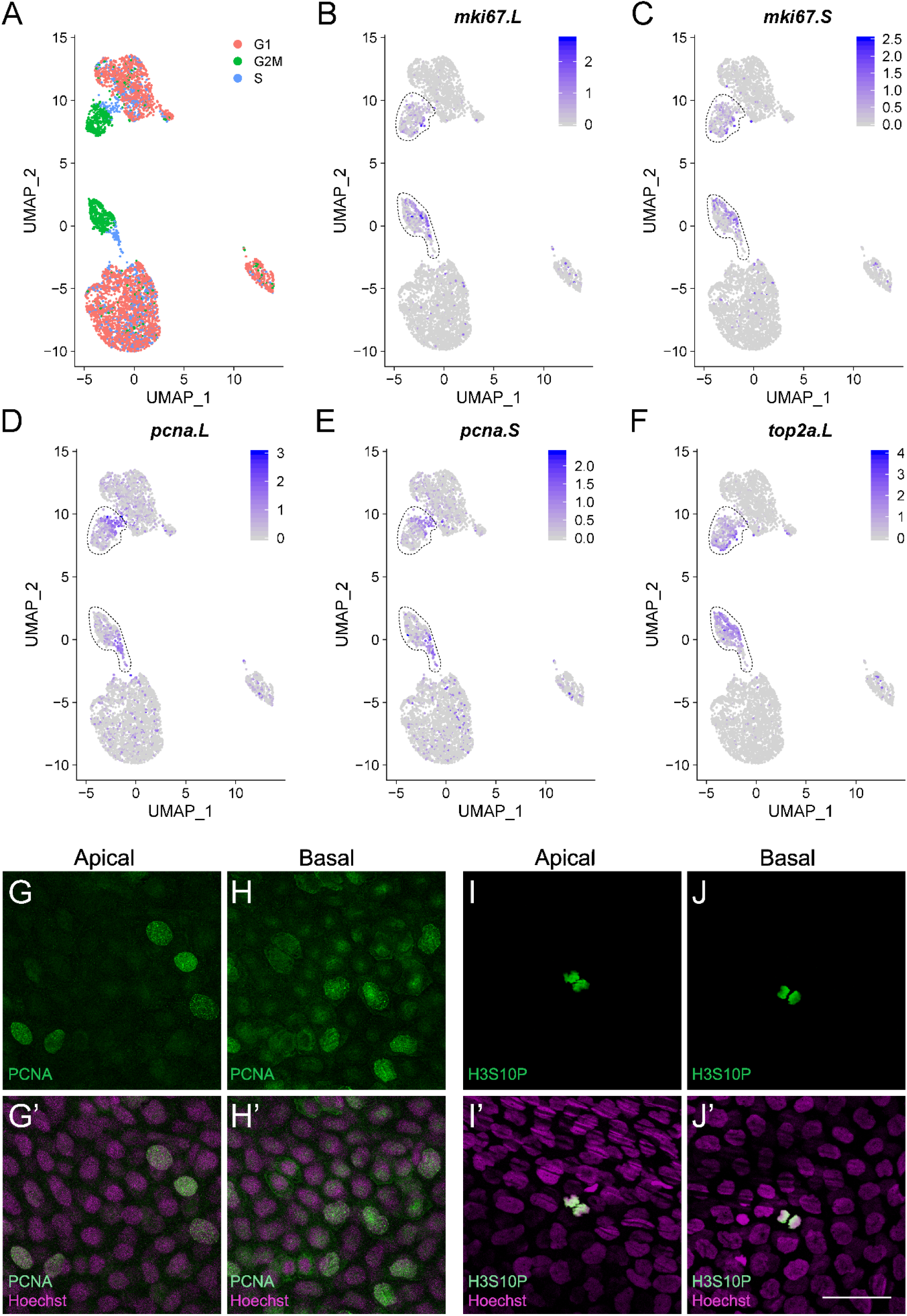
Proliferative cells reside in the bi-layered corneal epithelium of frog larvae. (**A**) UMAP plot showing the cell cycle status of each cell, as determined using the “CellCycleScoring” module in Seurat v3.1. The color key indicates the cell cycle state. (**B-F**) Gene expression UMAP overlay with proliferation marker genes *mki67.L*, *mki67.S*, *pcna.L*, *pcna.S* and *top2a.L*, respectively. Black dashed lines highlight clusters for the indicated gene. Color intensity correlates with the relative transcript level for the indicated gene in each cell, as noted in the key. (**G-H**) Confocal microscopic images showing immunofluorescent staining for Proliferating Cell Nuclear Antigen (PCNA) (green) in the larval frog corneal epithelium. (**G’-H’**) Merged images for G-H with Hoechst labeled nuclei (magenta). (G, G’) Nuclear staining for PCNA is detected in some apical epithelial cells. (H, H’) PCNA labeled nuclei are also detected in cells of the basal epithelium. (**I**-**J**) Confocal microscopic images showing immunofluorescent labeling for anti-phospho-Histone H3 (H3S10P) (green) in the tadpole corneal epithelium. (**I’**-**J’**) Merged images for I-J with Hoechst labeled nuclei (magenta). (I, I’) Mitotic nuclei (H3S10P) are present in a few apical epithelial cells. (J, J’) H3S10P labeled nuclei are also detected in a few cells of the basal epithelium. Scale bar in J’ equals 50 μm for G-J and G’-J’.

### Unique marker genes specify cluster identity in an unbiased manner

In organisms where detailed reference datasets are limited for select tissues and developmental stages, a notable advantage of scRNA-seq analysis is that it enables the identification of new marker genes and regulators of various cell types in an unbiased fashion. Genes with Differential Expression (DE) in a given cluster, when compared to all other clusters (average log fold change > 0.25; adjusted p value <0.01), were identified using Seurat’s “FindMarkers” function. Furthermore, the lists of DE genes were filtered by applying the criteria that cluster-specific unique marker genes be both highly and uniquely expressed. Therefore, we filtered DE genes using the difference between pct.1 and pct.2, along with a higher overall expression fold change value. For our analysis, we focused on genes expressed in more than 50% of the cells within that cluster. **Fig. 4A** depicts the heatmap with an expression profile of the top 10 candidate marker genes for each cluster, with the top 50 candidate marker genes being listed in **Table S3**. This unbiased approach identified genes with previously established roles in regulating other cell types and processes in *Xenopus*, as well as other vertebrates. Interestingly, as shown in **Fig. 4B**, the top-most marker gene in basal cells (Cluster 1) and apical cells (Cluster 4) is also shared with their proliferative cell clusters, namely clusters 2 and 3, respectively.

**Fig. 4.**
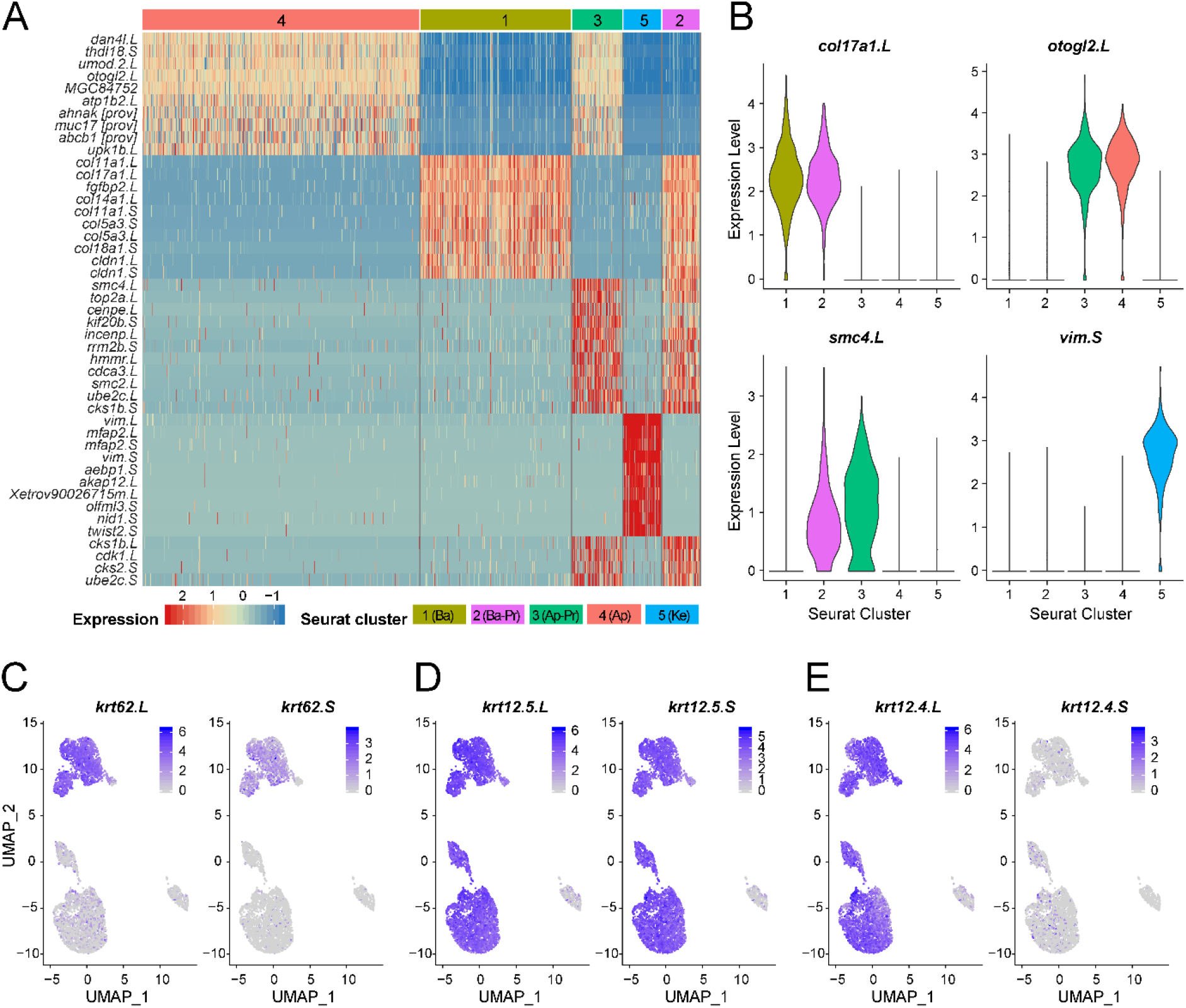
Identification of novel marker genes for different corneal cell types. (**A**) Heatmap showing the cluster-specific expression of the top 10 ranking novel candidate marker genes for each cell cluster identified in this scRNA-seq dataset. Relative expression and cluster identity are shown in the key. (**B**) Violin plots showing the cluster-specific expression of the top-ranking novel candidate marker genes for each identified cell type. Color and number correspond to the Seurat clusters shown in the key in (A). (**C-E**) Feature plots of .L and .S genes for *keratin 62* (*krt 62*), *keratin 12 gene 5* (*krt 12.5*) and *keratin 12 gene 4* (*krt12.4*), respectively. Relative expression is indicated in the color key.

Our examination of basal cells (Clusters 1 and 2; **Fig. 4A**) revealed that various collagen transcripts were significantly enriched in these cells, with *collagen type XVII alpha 1* (*col17a1.L)* being the most highly expressed marker gene (**Fig. 4B**). Collagens are the predominant proteins in the Extra Cellular Matrix (ECM) organization during corneal development and play diverse roles in cell adhesion and differentiation. Our discovery of *col17a1* transcripts in basal epithelial cells is particularly interesting. The gene *col17a1* encodes a hemidesmosomal transmembrane collagen protein, which is expressed in basal cells of the epidermis. A higher level of *col17a1* expression is associated as a marker of long-term epidermal stem cells and is crucial for skin homeostasis (Liu et al., 2019; Matsumura et al., 2016). That basal cells display an increased expression of *col17a1.L* points towards the possible role of this collagen gene as a unique marker and key regulator of CESCs and TACs present in this epithelial layer.

Looking at the apical corneal cells (Cluster 3 and 4; **Fig. 4A**), we detected the expression of larval skin-specific genes (e.g., *dan4l.L*) (Schreiber and Brown, 2003), goblet cell genes (e.g., *itln1*) (Dubaissi and Papalopulu, 2011), thyroid hormone-regulated genes (e.g., *thdl18.S*) (Furlow et al., 1997) and genes mediating signal transduction, as well as membrane permeability and stability (e.g., *upk1b.L*, *atp1b2.L*) (Eid and Brandli, 2001; Liao et al., 2018). The gene most uniquely enriched in apical cells encodes otogelin-like 2 (*otogl2.L*; **Fig. 4B**), a mucin glycoprotein secreted by small secretory cells (SSCs) and goblet cells of the larval epidermis. Otogl2 has been characterized as a major structural component of the mucosal protective barrier in *Xenopus* (Dubaissi et al., 2018; Dubaissi et al., 2014). Interestingly, in either of the apical epithelial clusters (Clusters 3 and 4) we did not detect the presence of genes associated with melanophores (*tyr*, *dct*, *mitf*, *sox10*) (Kawasaki et al., 2008; Kumasaka et al., 2003) that characterize the pigmented skin epithelium (**Fig. S4**). The absence of pigmentation is a characteristic feature of the transparent cornea, which otherwise is anatomically continuous with the surrounding skin.

We next focused on the proliferative cell clusters (Cluster 2 and 3) to identify their transcriptional heterogeneity. We determined that both these clusters had enriched expression of cell cycle-regulated genes (*incenp.L*, *cenpe.L*, *smc2.L*, *cdca3.L*; **Fig. 4A**), indicating that these cells are likely at different stages of cell cycle progression. The gene belonging to the Structural Maintenance of Chromosomes (SMC) family, *smc4.L* was the top marker gene in these cells (**Fig. 4B**). The SMC family member proteins are known to play crucial roles in mitotic chromosome condensation and segregation (Schmiesing et al., 2000). As one of the subunits of condensin I, SMC4 is widely conserved across eukaryotes and modulates chromosome structure for mitosis (Schmiesing et al., 1998).

For the corneal keratocyte cells (Cluster 5)— the neural crest-derived cells crucial for the formation and maintenance of the corneal stroma —a number of mesenchymal genes, particularly those involved in ECM deposition, cell adhesion and Epithelial to Mesenchymal Transition (EMT) (*twist2.S, mfap2.L*, *mfap2.S*, *nid1.S, olfml3.S*) were enriched (**Fig. 4A**). The top-most candidate in this gene list was vimentin (*vim.S*; **Fig. 4B**). As mentioned earlier, the presence of vimentin labeled keratocytes underneath the bi-layered epithelium has been described in tadpole corneas (Sonam et al., 2019). Overall, our dataset provides novel insights about a repertoire of genes that likely regulate the formation of corneal epithelial and stromal layers in these amphibians.

As various keratins play an integral role in the formation of the corneal epithelium, we also analyzed the cell-type-specific expression of some of the key keratins uniquely enriched in our dataset. Our results showed that the gene *keratin 62* (*krt62.L* and *krt62.S*; *krt62.L*, previously known as *xlk*) was highly abundant only in the basal corneal cells (Clusters 1 and 2; **Fig. 4C**). These findings concur with previous reports that this larval-specific keratin is solely expressed in basal cells of tadpole skin and its expression is turned off in the adult (Suzuki et al., 2009; Watanabe et al., 2001). Interestingly, two of the other larval epidermal keratins, namely *keratin 12 gene 5* (*krt12.5.L* and *krt12.5.S*, also known as *xk81b*) and *keratin 12 gene 4*, *.L* gene (*krt12.4.L*, also known as *xk81a1*) were enriched in apical as well as basal corneal cells, but not in corneal keratocytes. However, in the case of *krt12.4*, the *.S* gene had a much-reduced expression in apical and basal cells, suggesting subgenome-specific regulation and expression (**Fig. 4D-E**). This observation is consistent with earlier studies using *in situ* hybridization reporting the presence of *xk81* genes in the larval tail epidermis (Furlow et al., 1997).

### Claudin 4 and Keratin 8.2 differentially label the apical epithelial cells of tadpole cornea

To validate the spatial expression pattern for some of the newly identified genes, we performed immunohistochemical studies on corneal whole-mounts. This helped us confirm the cellular localization of genes, including *claudin 4* (*cldn4*) and *keratin 8* (*krt8.2*), which we determined as transcriptionally enriched in apical cell clusters (Cluster 3 and 4). Claudins are transmembrane proteins that form tight junctions, and several types of claudins are expressed in the human corneal epithelium. Among these, one of the most consistent and highly-expressed members is Claudin 4 (Ortiz-Melo et al., 2013; Yoshida et al., 2009). A recent study using whole mount preparations of post-metamorphic adult *Xenopus* corneas reported Cldn4 to be primarily localized to the lateral membranes of the surface corneal epithelium (Wiechmann et al., 2014). However, the authors did not examine Cldn4 expression in the corneal epithelium of younger tadpoles. In our results, we detected widespread Cldn4 immunolabeling in the membrane of the apical epithelial cells (Clusters 3 and 4, **Fig. 5A-B,B’**). However, the staining was absent in basal corneal cells (Clusters 1 and 2, **Fig. 5A,C,C’**), which agrees with our transcript dataset.

**Fig. 5.**
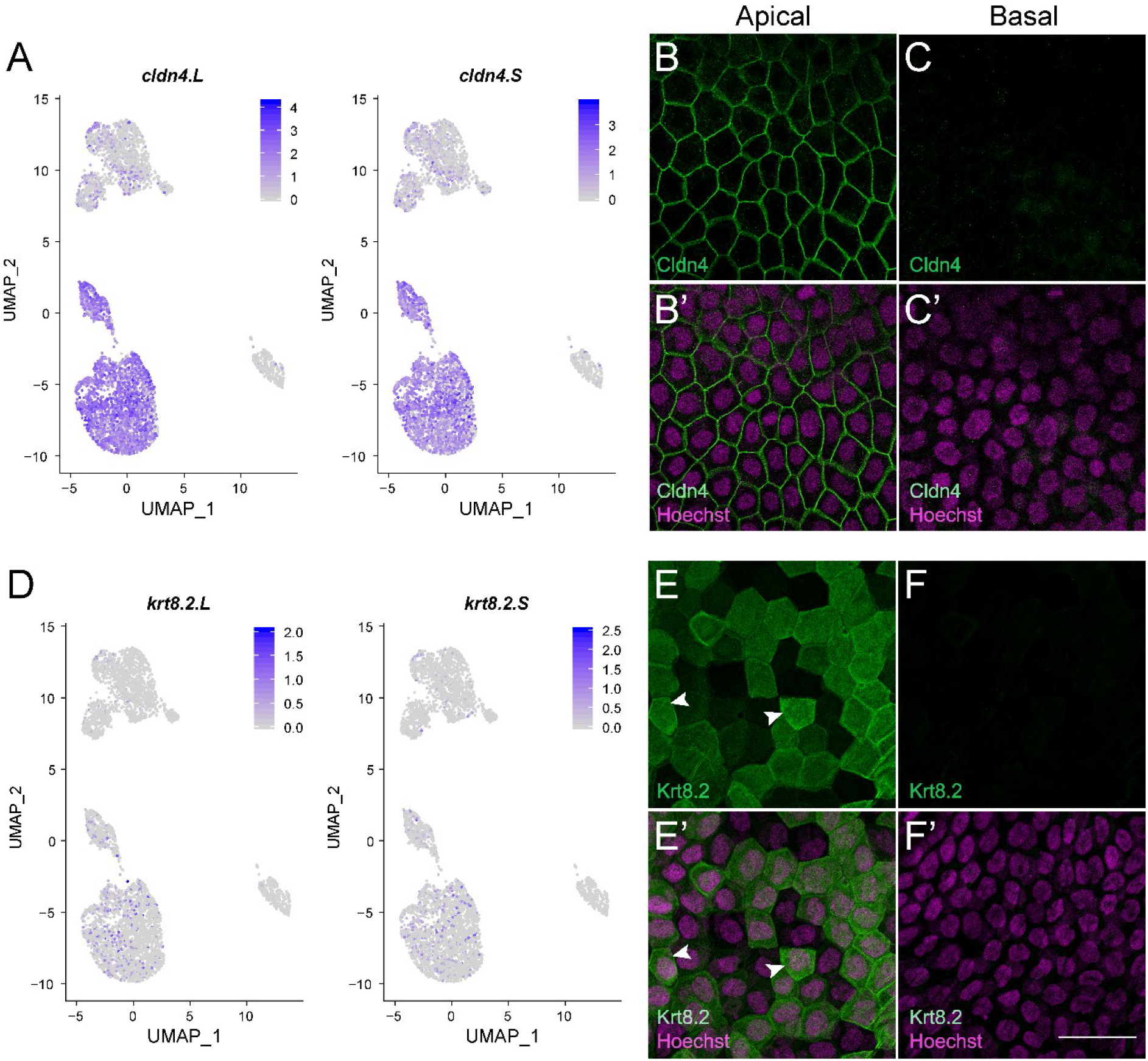
Validation of newly identified marker genes via protein localization in corneal whole mounts. (**A**) Feature plots depicting gene expression for *claudin 4* (*cldn4.L* and *cldn4.S*). Relative expression is indicated in the color key. (**B-C**) Confocal microscopic images showing immunofluorescent staining for Cldn4 (green) in the larval frog corneal epithelium. (**B’-C’**) Merged images for B-C with Hoechst labeled nuclei (magenta). (B, B’) Claudin 4 is uniformly expressed in the membranes of all epithelial cells of the apical layer. (C, C’) No label is detected in the basal epithelial cells. (**D**) Feature plots depicting gene expression for *keratin 8* (*krt8.2.L* and *krt8.2.S*). Relative expression is indicated in the color key. (**E-F**) Confocal microscopic images showing cellular localization of Krt8.2 (green) in the tadpole corneal epithelium. (**E’-F’**) Merged images for E-F with Hoechst labeled nuclei (magenta). (E, E’) Apical epithelial cells in the cornea show cytoplasmic staining for Krt8.2. A mosaic staining pattern is detected, with few cells displaying relatively higher expression than others (shown by white arrowheads). (F, F’) No Krt8.2 staining is present in basal corneal epithelial cells, which agrees with the transcript expression data. Scale bar in F’ equals 50 μm for B-C, B’-C’, E-F and E’-F’.

Another gene enriched in the apical cell clusters (Clusters 3 and 4) of the transcript dataset that we successfully validated was the intermediate filament protein, keratin 8.2 (*krt8.2.L* and *krt8.2.S*). Different type I and type II keratins are simultaneously expressed in keratinocytes—a major cell type of mammalian epidermis that maintains its barrier function (Ramms et al., 2013). Keratin 8, a marker of simple epithelium, is a prominent basic Type II keratin that is expressed early during epidermal development (Romano et al., 2012). A monoclonal *Xenopus* specific antibody (Kim et al., 2020; Mariani et al., 2020) was used to examine the corneal expression of this keratin. We observed a mosaic staining pattern, restricted to apical epithelial cells of the larval cornea (Clusters 3 and 4; **Fig. 5D-E,E’**). The expression was primarily cytoplasmic with some apical cells showing relatively higher expression than adjoining cells. The Krt8.2 staining pattern in apical cells may indicate that these cells are at different stages of cellular differentiation, as the cornea matures. On the other hand, no expression was detected in basal epithelial cells (**Fig. 5D,F,F’**). As reported in studies on mice epidermis formation (Romano et al., 2012), it is likely that the high nuclear expression of transcription factor p63 in basal cells (Sonam et al., 2019) is associated with suppressed expression of Keratin 8 in those cells.

### Analysis of gene expression trajectories reveals the differentiation pathway of corneal epithelial cells

As the vertebrate corneal epithelium is a self-renewing tissue (Kruse, 1994), it contains cells at different stages of differentiation (Beebe and Masters, 1996; Tseng, 1989). Therefore, to derive the differentiation trajectories of corneal epithelial cells, we leveraged one of the most powerful features of scRNA-seq datasets—single cells can be ordered in an unbiased way along a path to resolve their developmental progressions. We performed pseudo-time-based ordering of single cells from clusters 1, 2, 3 and 4 using Monocle’s network-based approach (Trapnell et al., 2014). The corneal keratocytes cluster (Cluster 5) was omitted from this analysis, as it was transcriptionally disconnected from the other four cell types (i.e., mesenchymal in origin). We found that the pseudo-time path has two branches (**Fig. 6A**), and different cell clusters can be arranged relatively clearly at different branch arms of this path. This pseudo-temporal ordering of cells along the axis originating from basal cells towards the terminally differentiated apical state was in agreement with the known corneal stratification in tadpoles (Hu et al., 2013). In principle, the distribution characteristics of different cell types on the differentiation trajectory can preliminarily describe the relationships between these cells during the development period. The distribution of basal cells (Cluster 1) and basal proliferative cells (Cluster 2) in the developmental trajectory is concentrated at the beginning (**Fig. 6B**), suggesting that their developmental relationships are close. Likewise, the apical epithelial cells (Cluster 4), along with the apical proliferative cells (Cluster 3) occupied a position around the second branch point, mostly towards the end of this path (**Fig. 6B**).

**Fig. 6.**
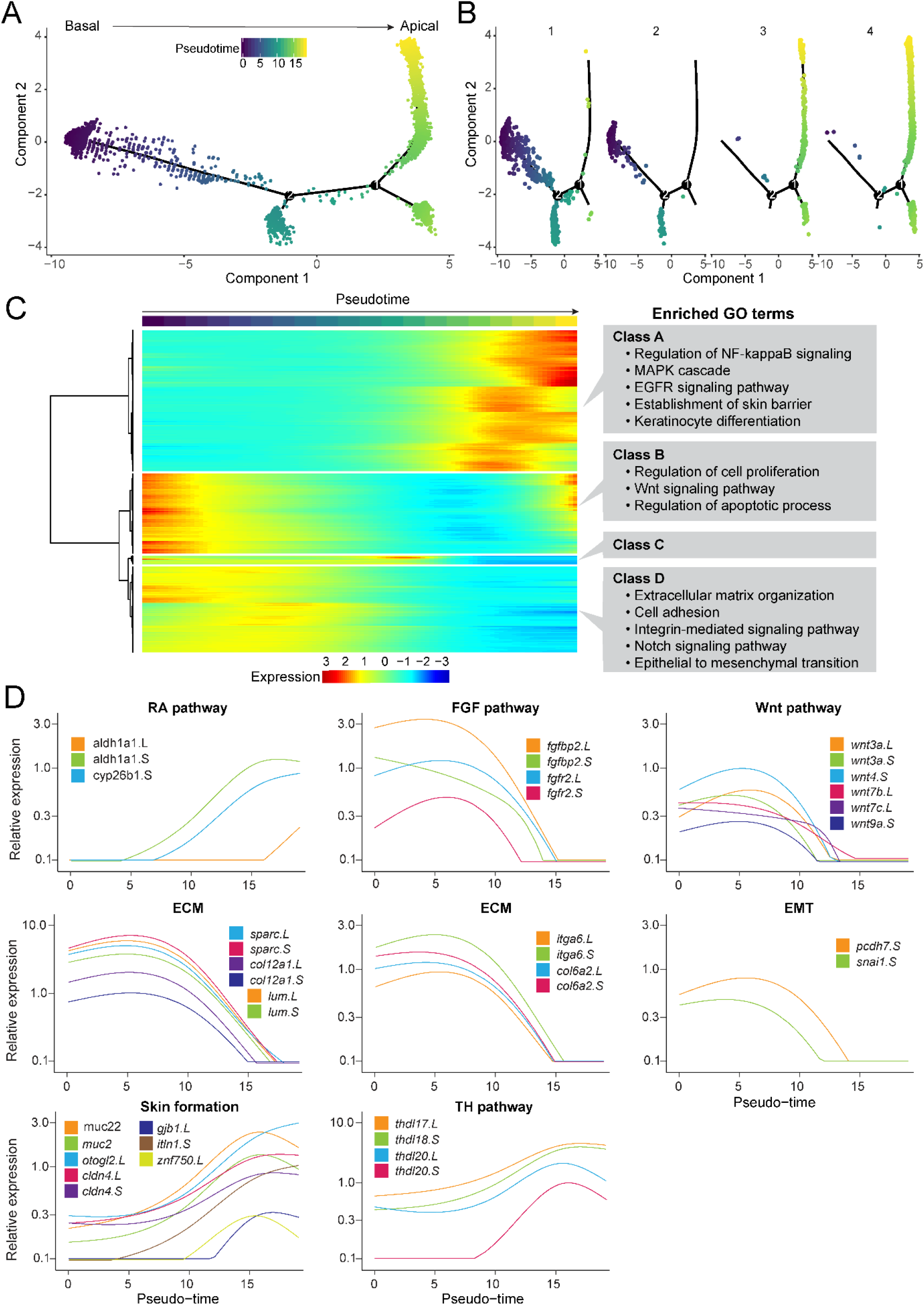
Pseudo-temporal ordering of individual cells reconstructs the corneal differentiation process. (**A**) Pseudo-time plot indicating the cellular trajectory of all corneal epithelial cells (Clusters 1-4), excluding the keratocytes. Single-cell trajectories were constructed, and pseudo-time values were calculated using Monocle 2. Trajectories are colored by pseudo-time, as indicated in the key. (**B**) Pseudo-time plots showing the distribution of each cell type along the combined cellular trajectories, as demonstrated in (A). Basal epithelial cells (Clusters 1 and 2) are present early along the trajectory, however, the apical cells (Clusters 3 and 4) are distributed more towards the end. (**C**) Heatmap depicting the classes of genes that vary along the pseudo-time during corneal epithelial differentiation. Pseudo-time is indicated by the color key similar to that shown in (A). Gene Ontology (GO) analysis was performed using DAVID and the enriched GO terms (p < 0.05) in each class are listed. Relative expression is indicated by the color key. (**D**) Gene expression kinetics along the pseudo-time progression of representative genes belonging to different pathways and processes, as indicated. Genes shown belong to the Retinoic Acid pathway (RA pathway), the Fibroblast Growth Factor pathway (FGF pathway), the Wnt pathway, Extracellular Matrix (ECM) deposition, Epithelial-to-Mesenchymal Transition (EMT), genes involved in Skin Formation and the Thyroid Hormone pathway (TH pathway).

Next, we identified the expression dynamics of the top 2000 genes that change as a function of progress through pseudo-time. These genes can be categorized into four distinct gene classes (namely Class A, B, C and D; **Fig. 6C**). To better understand the biological significance of these classes, we performed Gene Ontology (GO) analysis (see Material and Method). Genes predominantly expressed at the beginning of the developmental trajectory show an abundance of GO terms associated with cell proliferation, cell adhesion, integrin-mediated signaling, Wnt signaling, ECM organization and EMT (Classes B and D). At the other end of the trajectory, genes whose expression peaks late along this trajectory are enriched in GO terms associated with keratinocyte differentiation, the establishment of skin barrier functions and NF-kappaB signaling (Class A). In addition, the smallest class (Class C) with genes that peak transiently along the pseudo-time was enriched for ribosomal biogenesis and protein-membrane targeting genes (**Table S4**).

In addition to getting a coarse idea of broad GO terms that vary through corneal cell maturation, the pseudo-time trajectory can be used to gain insight into the temporal expression changes of individual genes. This provides us with a refined view of the changes that corneal epithelial cells undergo during their transition from basal to terminally differentiated apical states. As signaling mechanisms (such as Retinoic Acid, FGF and Wnt/β-catenin), processes of ECM deposition and EMT-related processes are crucial for corneal formation and differentiation (Dhouailly et al., 2014; Nakatsu et al., 2011), we closely investigated the expression changes of genes implicated in these pathways/processes along the pseudo-time (**Fig. 6D**). Retinoic acid signaling genes (*cyb26b1.S*, *aldh1a1.L*, *aldh1a1.S*) show an increasing trend along this pseudo-time, while genes in the FGF (*fgfbp2.L*, *fgfbp2.S*, *fgfr2.L*, *fgfr2.S*) and Wnt/β-catenin (*wnt3a.L*, *wnt3a.S*, *wnt4.S*, *wnt7b.L*, *wnt7c.L*, *wnt9a.S*) pathways show a reduction in expression. We also noted a dramatic reduction in expression of genes that primarily constitute the corneal ECM (*lum.L*, *lum.S*, *sparc.L*, *sparc.S*, *col12a1.L*, *col12a1.S*, *itga6.L*, *itga6.S*, *col6a2.L*, *col6a2.S*) along the progression of pseudo-time. A similar trend in reduction of expression was observed for EMT-related genes (*pcdh7.S*, *snai1.S*). On the other hand, genes characteristic of *Xenopus* skin formation (*otogl2.L*, *muc2, itln1.S* and *znf750.L*) gradually increase. Furthermore, we noted a remarkable increase in expression levels of thyroid hormone-controlled genes (*thdl18.S*, *thdl20.S* and *thdl17.L*) along the pseudo-time axis. Changes in thyroid hormone gene expression dynamics in frog cornea represents a novel observation and has not been characterized in earlier studies focused on *Xenopus* eye development. These changes are similar to those observed in frog skin (Suzuki et al., 2009; Yoshizato, 2007) and the corneal epithelium in these anurans likely also undergoes thyroid hormone-mediated remodeling—a hallmark characteristic of metamorphosis.

### Distinct gene regulatory networks are active during corneal differentiation

In order to characterize gene regulatory networks (GRNs) that govern corneal development and differentiation, we used the SCENIC pipeline to create a gene regulatory framework (Aibar et al., 2017). As transcription factors (TFs) are known to be highly conserved despite the allotetraploidization event in *Xenopus laevis* (Watanabe et al., 2017), we converted the *X. laevis* gene names to their human orthologs (see Materials and Methods). SCENIC calculates the activity of TFs from individual cells by integrating co-expression data with transcription factor motif enrichment analysis, to generate a “regulon.” A regulon is comprised of an expressed transcription factor and all of its co-expressed target genes. We determined the regulon activities using Area Under Cell (AUCell), which ranks targets of an individual regulon among the expressed genes, thereby generating a regulon-by-cell activity matrix.

To test our hypothesis that different developmental GRNs might drive cellular transitions during corneal differentiation, we analyzed the AUCell score activity matrix of individual corneal cells in our scRNA-seq dataset. Upon visualization of cell clustering from UMAP—built from the AUCell scores—we observed that corneal cells of a specific type grouped together, highlighting a high degree of similarity in their regulon activity (**Fig. 7A**). Importantly, the basal epithelial cells, including the proliferative population, formed distinct and non-overlapping clusters, which represent their discrete GRNs. Likewise, the proliferative cells of the apical epithelia clustered adjacent to apical cells and away from the basal population, demonstrating that two distinct regulons are active in the superficial corneal layer. The corneal keratocytes clustered distinctly away from both basal and apical cells, which indicates that regulatory networks driving these cells are strikingly different.

**Fig. 7.**
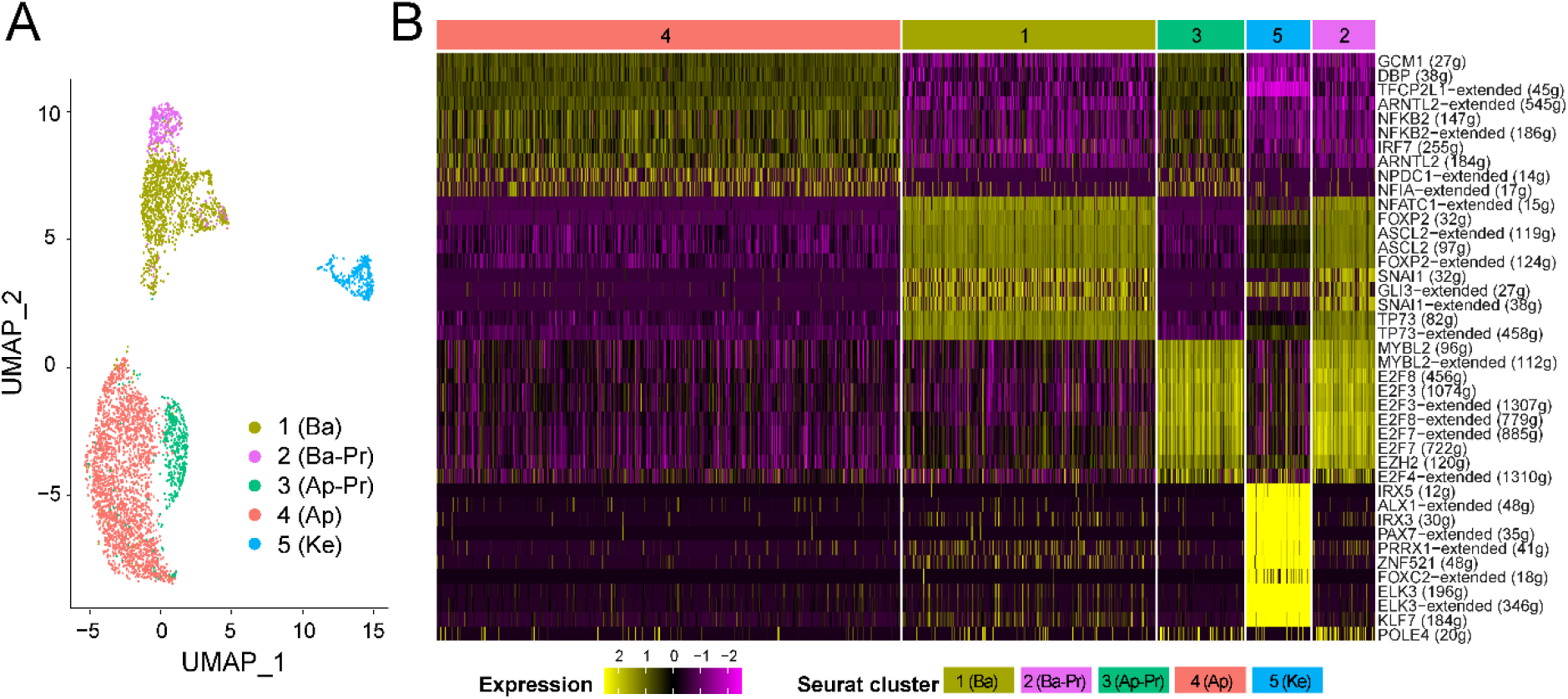
Distinct gene regulatory networks characterize the corneal cell types. (**A**) UMAP clustering of all corneal cells based on the AUC scores for each regulon calculated using SCENIC. Cells belonging to the same regulon and Seurat cluster are similarly colored, according to the key. (**B**) Heatmap depicting the different regulons that are active in different cell states. Relative expression is shown in the key.

In addition, we explored the regulons that are most upregulated in each of the cell clusters to identify the main TFs and their target genes that drive those cell states (**Fig. 7B**). In line with previous reports, in the outermost (apical) epithelial cells, we found increased expression of NF-kB, IRF7 and the ARNTL2 family of transcription factors that play essential roles in corneal wound healing, suppressing corneal infections and regulating corneal homeostasis (Alexander et al., 2006; Norman et al., 2004; Swamynathan, 2013). On the other hand, TP73, ASCL2, NFATC1, FOXP2 and SNAI1 regulons were active in basal cells and the basal proliferative cluster but were muted in apical clusters. As proteins of TF families such as the Forkhead box (e.g., FOXP2), basic helix-loop-helix (e.g., ASCL2) and Nuclear Factor of Activated T Cells (e.g., NFATC1) play a key role in maintaining a quiescent and undifferentiated state of stem cells (Keyes et al., 2013; Lay et al., 2016; Moriyama et al., 2008), these results support our previous hypothesis (Perry et al., 2013) that corneal stem cells reside throughout the basal epithelium in larvae. As expected, regulons associated with cell proliferation and progenitor maintenance such as E2Fs (E2F3,4,7,8) and EZH2 (Ezhkova et al., 2009; Francesconi et al., 2000) were enriched in the clusters of cycling cells. Regulons such as ELK3 (belonging to the ETS family), KLF7 (of the Kruppel-like Transcription factor family), PAX7 (of the Paired Domain-Homeodomain Transcription factor family) were highly active in corneal keratocytes, but had lowered expression in basal and apical epithelial cell clusters. Previous studies have described the role of these TF families in regulating cell differentiation, cell migration, and matrix formation processes during corneal development (Klein et al., 2017; Yoshida et al., 2000).

## DISCUSSION

Here, we have used single-cell transcriptional profiling to probe cellular heterogeneity in corneal epithelial tissue of *Xenopus* larvae. Although the study captures a narrow time-point of larval corneal development, it enabled us to generate a high-resolution atlas that describes the major cell types and reconstructs developmental trajectories of corneal cells under homeostatic conditions. A comprehensive characterization of cell-type-specific marker genes along with an understanding of the corneal epithelial developmental lineage and dynamic regulatory network of key transcription factors will provide future avenues to optimize the maintenance and differentiation of corneal cells in the frog and other vertebrates.

### The larval cornea comprises five discrete cell types, including proliferative cells in both apical and basal layers

The formation of the *Xenopus* larval epidermis has been extensively characterized in several studies and remains of critical biomedical interest (Boon et al., 2014; Walentek and Quigley, 2017). Nevertheless, despite sharing a high degree of developmental and anatomical similarity to human corneas (Hu et al., 2013), the gene expression profile of frog corneal development has not been previously characterized. Our study profiled nearly five thousand corneal cells, and this data offers valuable insights into cellular processes that characterize distinct corneal cell types—basal cells (Cluster 1), apical cells (Cluster 4), corneal keratocytes (Cluster 5) and proliferative cells of basal and apical epithelia (Clusters 2 and 3, respectively). The identification of proliferative cells contained in both the epithelial layers was intriguing (**Fig. 3**), as past studies have postulated that proliferative cells exist mainly within the basal epithelial layer and upon differentiation, these cells populate the outermost layer of the corneal epithelium (Li et al., 2017; Perry et al., 2013; Sun and Lavker, 2004; Yoon et al., 2014).

Using an unbiased approach to marker gene calling, we identified expression features that define each cluster and determined unique genes that characterize and regulate these cell states (**Fig. 4**). Examination of specific corneal layers revealed that the outer (apical) layer of corneal epithelium expresses genes associated with specialized epidermal cell types (goblet cells, ionocytes, small secretory cells) but lacks pigmented melanophore genes. The absence of pigmentation is expected in the cornea, as maintaining optical clarity is essential for proper vision. The inner (basal) epithelial layer is characterized by an abundance of ECM genes, including various collagens, integrins, and keratins. Some of the newly identified basal marker genes (e.g., *col17a1*, *col5a3*, *cldn1*, *krt62* and *smc4*) may contribute as regulators of corneal epithelial development and differentiation, particularly of CESCs and early TACs, in this amphibian and other vertebrates.

### Identification of dual populations of quiescent and proliferative cells

Analyses of the data reveal two distinct populations of proliferative cells: one that is closely related to those of the quiescent basal cell cluster and another that is closely associated with cells of the quiescent apical cell cluster. The basal proliferative cells (Cluster 2) showed an abundance of basal cell markers, also expressed in the non-proliferative basal cells of Cluster 1 (e.g., *col17a1.L*, *krt19.L*). In contrast, the apical proliferative cells (Cluster 3) expressed differentiated apical cell markers found in the non-proliferative apical cells of Cluster 4 (e.g., *cldn4.L*, *otogl2.L*) (see **Fig. 4** and **Table S3**).

Previous studies considered genes such as *Mki67* and *PCNA* to represent potential markers for proliferative TACs (Guzman et al., 2013; Lehrer et al., 1998). In fact, both proliferative cell clusters (Clusters 2 and 3) exhibited an enriched expression of these proliferation marker genes. These populations of basal and apical proliferative cells, respectively, could represent the so-called “early TACs” and “late/mature TACs” proposed in a recent scRNA-seq study of the mice corneal epithelium (Kaplan et al., 2019). However, we could not compare the entire corneal epithelial gene expression profile of mouse to that of tadpoles, due to unavailability of raw data for the mice study.

Based on the larval corneal expression profiles, it is still a challenge to discern stem cells from early stage TACs—a conclusion in line with other studies focused on identifying specific markers for CESCs and TACs (Lavker et al., 2020; Mort et al., 2012). In fact, it is possible that the two basal corneal epithelial cell populations (Clusters 1 and 2) represent only one cell type, and our analysis has simply captured those cells when they are also expressing genes that regulate mitosis.

While we did not identify a smaller discrete population of CESCs, we did observe a higher level of expression of epithelial stem-cell related marker genes (e.g., *tp63, krt19*, *krt15.2*, *cldn1, itga6*, *itgb4*, *col17a1*) in cells of both basal clusters (Clusters 1 and 2). The similar transcriptional profiles of these clusters suggest that the entire basal cell population behaves like stem cells in *Xenopus* larvae. This observation supports an earlier proposal that oligopotent stem cells may be distributed throughout the basal corneal epithelium during larval stages (Perry et al., 2013). In future studies, it will be important to examine the adult frog cornea, where evidence suggests that EdU label-retaining putative CESCs represent a relatively smaller population of cells that reside in the limbus (Hamilton and Henry, 2016). The altered spatial distribution of cells in the mature cornea may affect gene expression dynamics, and perhaps one will be able to better distinguish CESCs and TACs.

It is also of some significance that we did not find evidence for a more gradual transition in gene expression, as the corneal epithelial cells undergo differentiation (and a greater number of transitional cell types/clusters). Instead, our observations suggest a model in which a more discrete genetic switch occurs as cells transition from basal to apical fates.

### Variable developmental trajectories and discrete TF regulatory networks govern the process of corneal maturation

To dissect the temporal and spatial distribution of corneal cells, we performed a pseudo-time analysis of our scRNA-seq data (**Fig. 6**). As illustrated by the waves of gene expression identified here, cells transitioning from basal to apical layers follow finely resolved developmental trajectories. Our study offers the required resolution to distinguish the stepwise temporal progressions connecting the beginning and end of corneal epithelial differentiation. Although it cannot be ruled out that the progressions that individual cells undergo during differentiation may be more dynamic than captured in this analysis, this data definitely adds to our current understanding of corneal development and differentiation. We are particularly intrigued by our novel finding of the role of thyroid hormone genes in corneal differentiation, and seek to explore this in future studies. Furthermore, a number of signaling pathways appear to be involved (e.g., Retinoic Acid, FGF, Wnt/β-catenin, BMP) that have also been shown to play crucial roles in supporting cornea-lens regeneration in frogs (Day and Beck, 2011; Fukui and Henry, 2011; Hamilton et al., 2016; Thomas and Henry, 2014). In the future, it will be interesting to examine the gene expression profiles and developmental trajectory of regenerating individual corneal cells across various stages of lens reformation and discern their spatiotemporal progressions.

At the transcriptomic level, the expression patterns of many key genes involved in corneal epithelial development are known, but regulators of these genes are still being identified (Stephens et al., 2013; Swamynathan, 2013). Here, we leveraged our scRNA-seq data to computationally infer dynamic TF networks controlling corneal cell states, as well as its maturation process. Distinct groups of TFs characterize cell fate changes during the progression from the undifferentiated basal to the differentiated apical states (**Fig. 7**). Many of these newly identified TFs and their target genes (e.g., FOX and bHLH for basal cells and NF-kB and IRF for apical cells) can be explored further as key regulators of corneal epithelium formation and maintenance, including that of CESCs and TACs in vertebrates.

### Future scRNA-seq to elucidate cornea-lens regeneration in frogs

Despite sharing an extensive parallel with skin epithelium both in terms of cell types and transcriptional profile, the corneal epithelium in these anurans is unique. Only the tadpole cornea and pericorneal epithelium (lentogenic area—a combined region extending twice the diameter of the eye) is capable of regenerating a functional lens. On the other hand, the flank epidermis (outside this lentogenic area) does not possess this competence (Cannata et al., 2003; Freeman, 1963). Earlier work on *Xenopus* lens regeneration had identified some candidate genes, including *pax6* and *fgfr2* (*bek isoform*), that may contribute to the cornea’s capacity to regrow a lens (Arresta et al., 2005; Gargioli et al., 2008). Future scRNA-seq studies of tadpole flank epidermis will enable us to compare the molecular heterogeneity of both these epithelia, to identify the entire gene repertoire that contributes to the cornea’s capacity to regenerate a lens. Likewise, it will be valuable to examine the corneal epithelium of metamorphosed frogs to understand what transcriptional changes lead to its lost lens regenerative capacity (Filoni et al., 1997; Hamilton and Henry, 2016).

### *Xenopus laevis* as a corneal disease model

*Xenopus laevis* is a well-established model for studying eye development and regeneration processes (Henry et al., 2008). These frogs share a high degree of synteny with humans, with around 80% of human disease genes having orthologs in these amphibians—making them an excellent choice to model human diseases (Blum and Ott, 2018; Nenni et al., 2019). Recently, robust *Xenopus* ocular disease models have been established for congenital eye malformations including aniridia, cataract, megalocornea, microphthalmia, coloboma and corneal stem cell deficiency (Adil et al., 2019; Nakayama et al., 2015; Pfirrmann et al., 2015; Viet et al., 2020). Along with being easily accessible for experimental manipulation and imaging, the frog cornea resembles that of humans (Hu et al., 2013) that makes it a suitable vertebrate model for human corneal disease-related investigations. In our current study, several genes associated with human corneal dystrophies were found to be expressed in the tadpole cornea (e.g., *LUM*, *SLC4A11*, *SOD1* and *KERA*; see **Fig. S5**)(Klintworth, 2009). For instance, the putative *Xenopus* ortholog of the *KERATIN 12* (*KRT12*) gene related to Meesmann’s corneal dystrophy (Irvine et al., 2002; Irvine et al., 1997) was expressed significantly in cells of the apical and basal epithelium (i.e., *krt12.5.L*, *krt12.5.S*, *krt12.4.L*; **Fig. 4D-E**). With the advanced genetic manipulation tools now available for *Xenopus* (Tandon et al., 2017), frogs can be developed to model human corneal dystrophies and to better understand disease pathobiology.

## Conclusion

In summary, the transcriptional atlas of the *Xenopus* cornea described here provides a unique spatiotemporal perspective of corneal epithelium differentiation. One can extrapolate the findings of this study to test a number of newly identified enriched basal corneal epithelial genes for their potential application as specific markers in other vertebrates, including mice and humans. This also indicates that scRNA-seq can be used as a sensitive approach to comprehensively document transcriptional profiles at various stages of mammalian corneal development to reconstruct precise developmental lineages of corneal cells. An extensive molecular and cellular characterization of corneal epithelia among various vertebrates will ultimately aid in determining accurate biomarkers for the identification and isolation of stem cells for clinical purposes. This will also be crucial for designing and improving therapeutic treatments for diseases and dystrophies that affect corneal epithelial integrity.

## MATERIALS AND METHODS

### Animals

Adult *Xenopus laevis* were obtained from Nasco (Fort Atkinson, WI). Fertilized eggs were collected, and larvae were raised to Stage 49-51 based on developmental staging by Nieuwkoop and Faber (1956). All animal care and experiments performed in this study were approved and monitored by the Institutional Animal Care and Use Committee (IACUC) and the Division of Animal Resources (DAR) at the University of Illinois.

### Single-cell dissociation

Tadpoles were anesthetized at room temperature by incubation for 30-60 seconds in a 1:2000 dilution of MS222 (ethyl 3-aminobenzoate methanesulfonate, Sigma, St. Louis, MO) diluted in 1/20X Normal Amphibian Media, NAM (Slack, 1984). Anesthetized tadpoles were transferred to clay-lined dishes containing Ca^++^−Mg^++^ free Dulbecco’s Phosphate Buffered Saline (1X DPBS; Corning, NY, #21-031-CV). The corneal epithelial tissue was dissected using fine microscissors (Henry et al., 2018), taking care not to include any surrounding skin. The skin is demarked by the outer edge of the orbit, a lack of transparency, and the obvious presence of pigmented melanophores. To limit the amount of time needed to collect these tissues, three operators simultaneously collected a total of approximately 80 corneas in three separate glass depression dishes containing 400 μl 1X DPBS, on ice. Care was taken to ensure that corneas were carefully removed from the tip of the forceps and that they remained submerged in the 1X DPBS solution. Using a 200 μl pipette tip, most of the DPBS was gently removed without disturbing the corneas at the bottom of each glass dish. To dissociate the corneal epithelial tissue, ~350 μl of Accutase (CellnTec Advanced Cell Systems, Bern, Switzerland) was added to each dish, and the dishes were covered with plastic petri dish lids (35 × 10 mm, Corning, #351008) flat side down, and wrapped tightly with lab tape to secure them and prevent desiccation. Next, the dishes were incubated at 37°C for 35 minutes with intermittent trituration using a 200 μl pipette tip. To stop the reaction, 200 μl of media containing 61% L-15 (Invitrogen, Carlsbad, CA), 0.5 μM EDTA (Invitrogen) and 10% Fetal Bovine Serum (Invitrogen) was added to each dish. For mechanical dissociation, the suspension was finally triturated 8-10 times using the 200 μl pipette tip. The three cell suspensions were pooled in a 2ml LoBind microcentrifuge tube (Eppendorf, Enfield, CT; #022431048), spun at 325g for 5 minutes in a fixed angle centrifuge, and resuspended in 200 μl L-15 with 10% FBS and 0.5 μM EDTA. Finally, the cells were passed through a 40 μm FlowMi cell strainer (Bel-Art, Wayne, NJ; #136800045) and collected in a 2ml LoBind tube. Cell viability was assessed using Acridine Orange/Propidium Iodide (AO/PI) staining solution (Nexcelom Bioscience, Lawrence, MA, #CS2-0106) that labels live and dead cells respectively, and counted using an automated cell counter (Cellometer K2, Nexcelom Bioscience).

### Library preparation and sequencing

The single-cell RNA-seq library was generated using the 10X Genomics Chromium Single Cell 3’ Kit v3 (10X Genomics, San Francisco, CA) and sequenced with Illumina NovaSeq 6000 on a SP flow cell to obtain 150bp paired reads. Library preparation and sequencing was conducted at the High-Throughput Sequencing and Genotyping Unit of the Roy J. Carver Biotechnology Center at the University of Illinois.

### scRNA-seq data processing

RNA sequencing reads for the 10X library were processed using the CellRanger v3.1.0 pipeline by 10X Genomics. For generating the CellRanger reference, the *Xenopus laevis* v9.2 reference genome and gene models were downloaded from Xenbase (http://ftp.xenbase.org/pub/Genomics/JGI/Xenla9.2/XENLA_9.2_Xenbase.gff3) (Karimi et al., 2018). Further downstream processing was performed in R Studio v3.6.1. The output from CellRanger was uploaded into an R Studio session, and processed using the Seurat v3.1 pipeline (Butler et al., 2018; Satija et al., 2015), wherein a standard filtering and preprocessing pipeline was employed. Genes expressed in less than 3 cells, and cells expressing <200 or >3000 genes were removed from downstream analysis. We used a 5% cutoff for mitochondrial reads to remove dead/apoptotic cells. Post-normalization, we scaled data using the ScaleData function and performed this for all features/genes. We used FindNeighbours and FindClusters functions at parameter values of dims 1:20 and resolution 0.1, respectively. Significantly enriched genes within clusters were identified using the FindMarkers function, with a min.pct parameter of 0.25. Cell type identities were assigned using expression patterns of previously established marker genes from the available literature (Castro-Muñozledo, 2015; Guo et al., 2018; Li et al., 2017) and our previous study (Sonam et al., 2019). The frog *X. laevis* is allotetraploid, and gene expression data can be accessed for each allele from both the Large (Gene.L) and/or Short (Gene.S) chromosomes (Session et al., 2016). Taking into consideration that these alleles may have different functions, we reported cluster-specific allele expression patterns for each gene.

### Trajectory Analysis

To infer the trajectory of single cells, we used Monocle v2.0 (Trapnell et al., 2014) and applied the DDRtree algorithm for dimensionality reduction. Differentially expressed genes were calculated using comparison across the different clusters found by Seurat v3.1 above. We used the Top 3000 genes to order cells and determine the pseudo-time trajectory. Relevant functions within Monocle2 were used to plot a heatmap for genes varying along the pseudo timeline. To define the Gene Ontology (GO) for the regulated genes along the trajectory, we humanized the *X. laevis* gene names and converted them to human Ensembl GeneIDs, and used DAVID 6.8 for the GO analysis (Dennis et al., 2003). GO option GOTERM_BP_ALL was selected, and the terms with a p-value < 0.05 were chosen as significant categories.

### Cell Cycle Analysis

For cell cycle analysis, we used a previously defined core set of 84 G1/S and 98 G2/M cell cycle genes (Aztekin et al., 2019). We identified the approximate cell cycle state of each cell with the average expression (called the “cycle score”) for the two gene sets using the Cell cycle scoring function in Seurat v3.1. If the cycle scores of G1/S and G2/M were both less than 2, the corresponding cell was considered to be quiescent; otherwise, the corresponding cell was identified as being proliferative.

### Gene Regulatory Network (GRN) analysis

We used the single-cell regulatory network inference and clustering (SCENIC) pipeline (Aibar et al., 2017) to estimate the AUCell Score activity matrix from the scRNA-seq dataset (Chembazhi et al., 2020). Briefly, since current relevant databases don’t hold gene regulatory information specific to *X. laevis*, we decided to collapse raw counts from .L and .S gene forms in our data. This collapsed data was then processed in the Seurat v3.1 pipeline, as explained above. A normalized count table was extracted from the Seurat object, and *Xenopus* genes with a known human ortholog were retained for downstream analysis, and the gene names were humanized. Unlike the standard SCENIC workflow where this AUCell score activity matrix is binarized by thresholding to generate binary regulon-activity matrix, we retained the full AUCell score for all further analysis. The matrix of AUCell score containing regulons (rows) x cells (columns) was used as input to Seurat object and used to cluster and identify important regulons for each previously identified cluster.

### Immunofluorescence staining and imaging

Immunostaining of larval corneas was performed using a previously published protocol (Sonam et al., 2019). Briefly, the larval specimens were fixed in Dent’s fixative (80% methanol, 20% dimethyl sulfoxide, DMSO), or 3.7% ParaFormaldehyde (PFA) fixative. Next, the eyes were excised from each sample, taking care to keep the cornea intact. The excised eyes were washed six times for 10 minutes each in 1X PBST (Phosphate buffered saline; 1.86 mM NaH_2_PO_4_, 8.41 mM Na_2_HPO_4_, 175 mM NaCl, pH7.4, containing 0.5% Tween-20). Blocking was done using 5% Bovine Serum Albumin/10% normal goat serum/1X PBST for 2 hours at room temperature. Finally, the eyes were incubated with primary antibodies diluted in the blocking solution overnight at 4°C (see **Table S1**). Following primary incubation, six more washes were done with 1X PBST prior to incubating the eyes with secondary antibodies. Goat anti-mouse Alexa-Fluor 488 or 546, and goat anti-rabbit Alexa-Fluor 488 or 546 (Invitrogen, Rockford, IL) were used as secondary antibodies at a concentration of 1:300 diluted in blocking solution for 2 hours at room temperature. Finally, the eyes were washed twice with 1X PBST and incubated with 1μg/ml solution of Hoechst 33342 (Molecular Probes, Eugene, OR) in 1X PBS for nuclear counter-staining. RainX (ITW Global Brands, Houston, TX) treated slides were used for mounting the corneas, and SlowFade antifade reagent (Life Technologies, Eugene, OR) was used for the mounting media. To flat mount the cornea on a slide, labeled corneas were carefully removed from the whole eyes, and small, peripheral, radial incisions were made to facilitate flattening during the mounting process.

Confocal imaging of immunolabeled larval corneas was performed using an inverted LSM 700 microscope (Carl Zeiss, Munich, Germany). The Z-stacks were processed with Zen software (Carl Zeiss, Munich, Germany). The images were compiled using Adobe Photoshop.

## ACKNOWLEDGEMENTS

We acknowledge support from the High-throughput sequencing center, UIUC and the Developmental Studies Hybridoma Bank for 1H5 antibody.

## DATA AVAILABILITY

All raw RNA-seq data files are available for download from the NCBI Gene Expression Omnibus (http://www.ncbi.nlm.nih.gov/geo/) under accession number GSE154896.

## SUPPLEMENTARY FIGURES

**Fig. S1.**
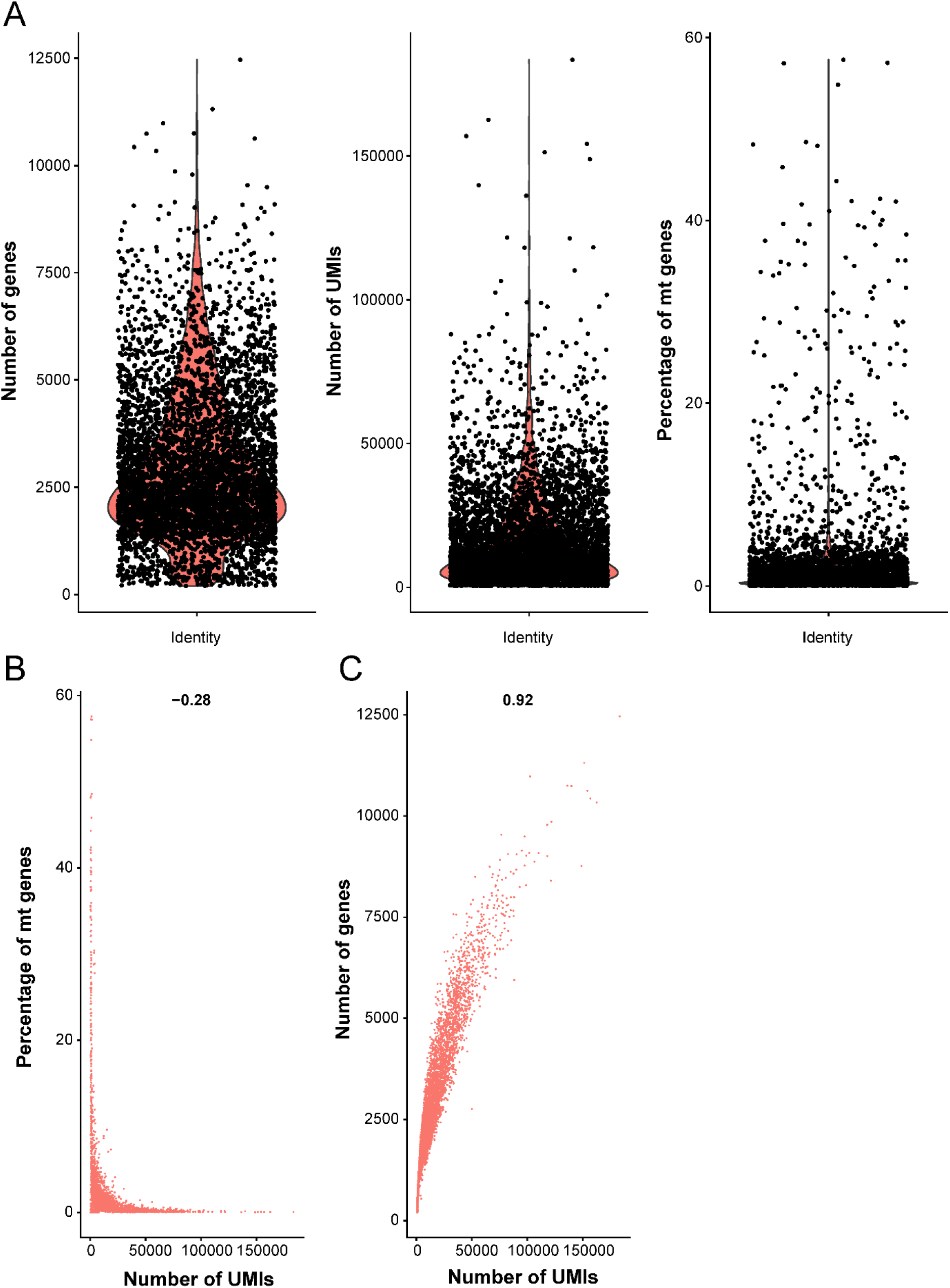
Quality control (QC) metrics of *Xenopus* larvae corneal epithelial scRNA-seq data. (**A**) Violin plot illustrating the distribution of number of genes, unique molecular identifiers (UMIs) and the percentage of mitochondrial (mt) genes, respectively, in each cell (black dot) of the pooled corneal sample. (**B**) Scatterplot showing the correlation of percentage of mitochondrial genes and UMI count. (**C**) Scatterplot showing the correlation of number of genes and number of UMIs.

**Fig. S2.**
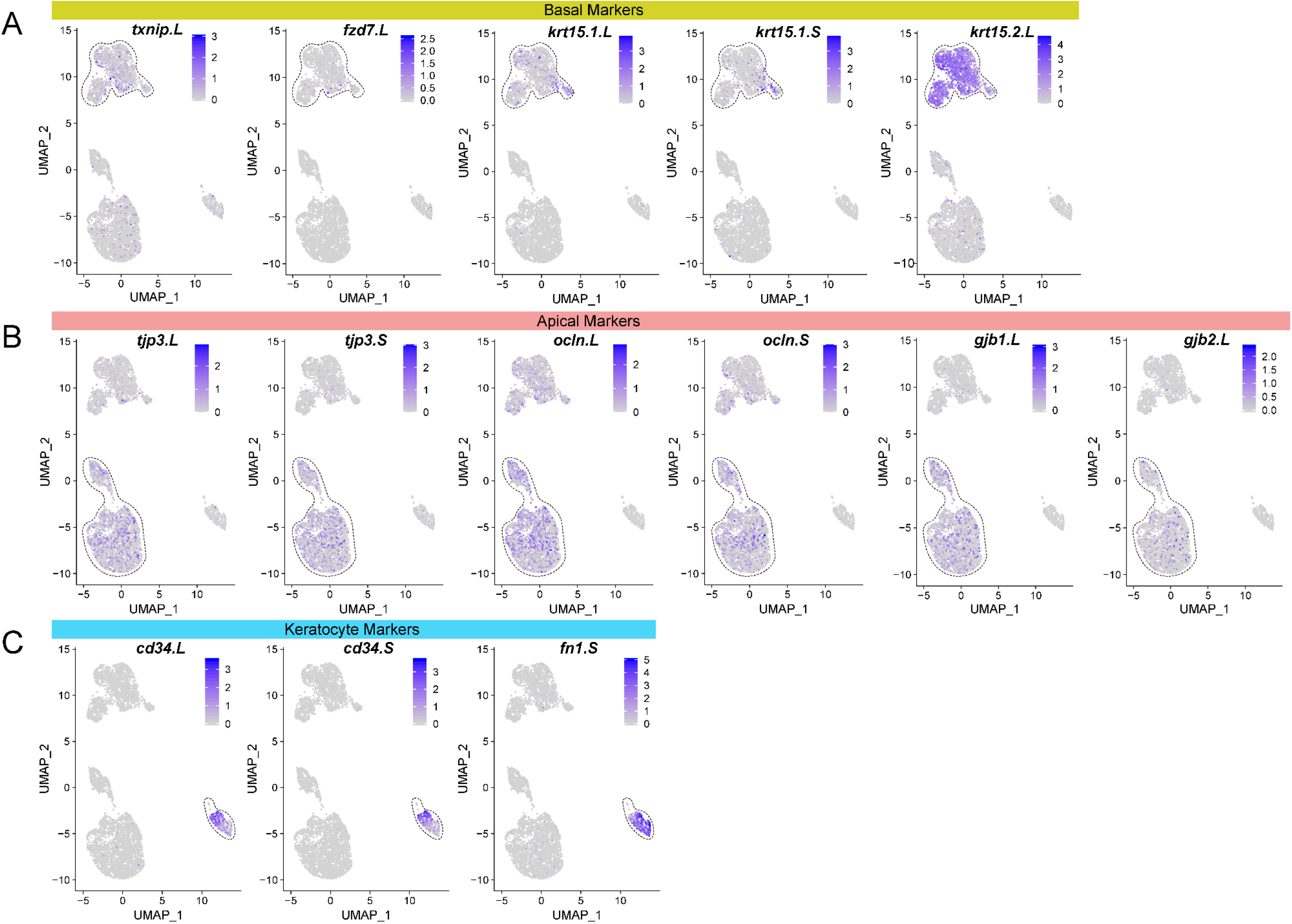
(related to Fig. 2) Cell cluster identity was further validated based on the expression of known marker genes. Feature plots of major cell-type-specific marker genes, known from the literature (see text). Black dashed lines highlight the cluster where the indicated gene was more highly expressed. (**A**) Corneal basal cell marker genes: *thioredoxin interacting protein* (*txnip.L*); *frizzled class receptor 7* (*fzd7.L*); *keratin 15 gene 1* (*krt15.1.L* and *krt15.1.S*) and *keratin 15 gene 2* (*krt15.2.L*). (**B**) Apical epithelial cell marker genes: *tight junction protein 3* (*tjp3.L* and *tjp3.S*); *occludin* (*ocln.L* and *ocln.S*); *gap junction protein beta 1* (*gjb1.L*) and *gap junction protein beta 2* (*gjb2.L*). (**C**) Keratocyte marker genes: *CD34 molecule* (*cd34.L* and *cd34.S*) and *fibronectin 1* (*fn1.S*). Relative expression is indicated in the keys.

**Fig. S3.**
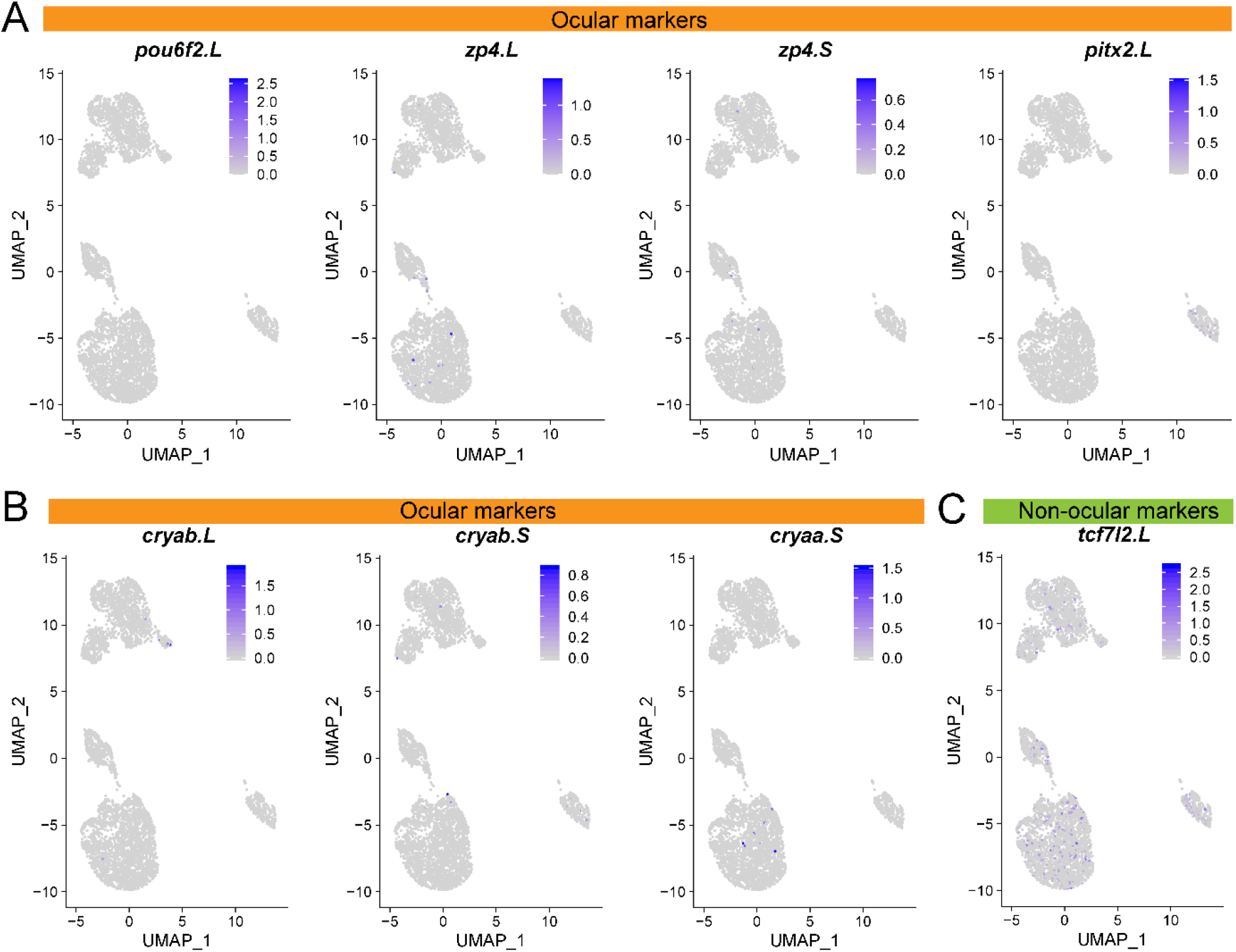
The absence of other ocular and non-ocular cell types was verified based on the expression of marker genes. Feature plots of cell-type-specific marker genes. (**A**) Corneal endothelial marker genes: *POU class 6 homeobox 2* (*pou6f2.L*); *zona pellucida glycoprotein 4* (*zp4.L* and *zp4.S*) and *paired like homeodomain 2* (*pitx2.L*). (**B**) Lens cell marker genes: *crystallin alpha B* (*cryab.L* and *cryab.S*) and *crystallin alpha A* (*cryaa.S*). (**C**) Marker gene to differentiate cornea versus skin: *transcription factor 7 like 2* (*tcf7l2.L*). Relative expression is indicated in the keys.

**Fig. S4.**
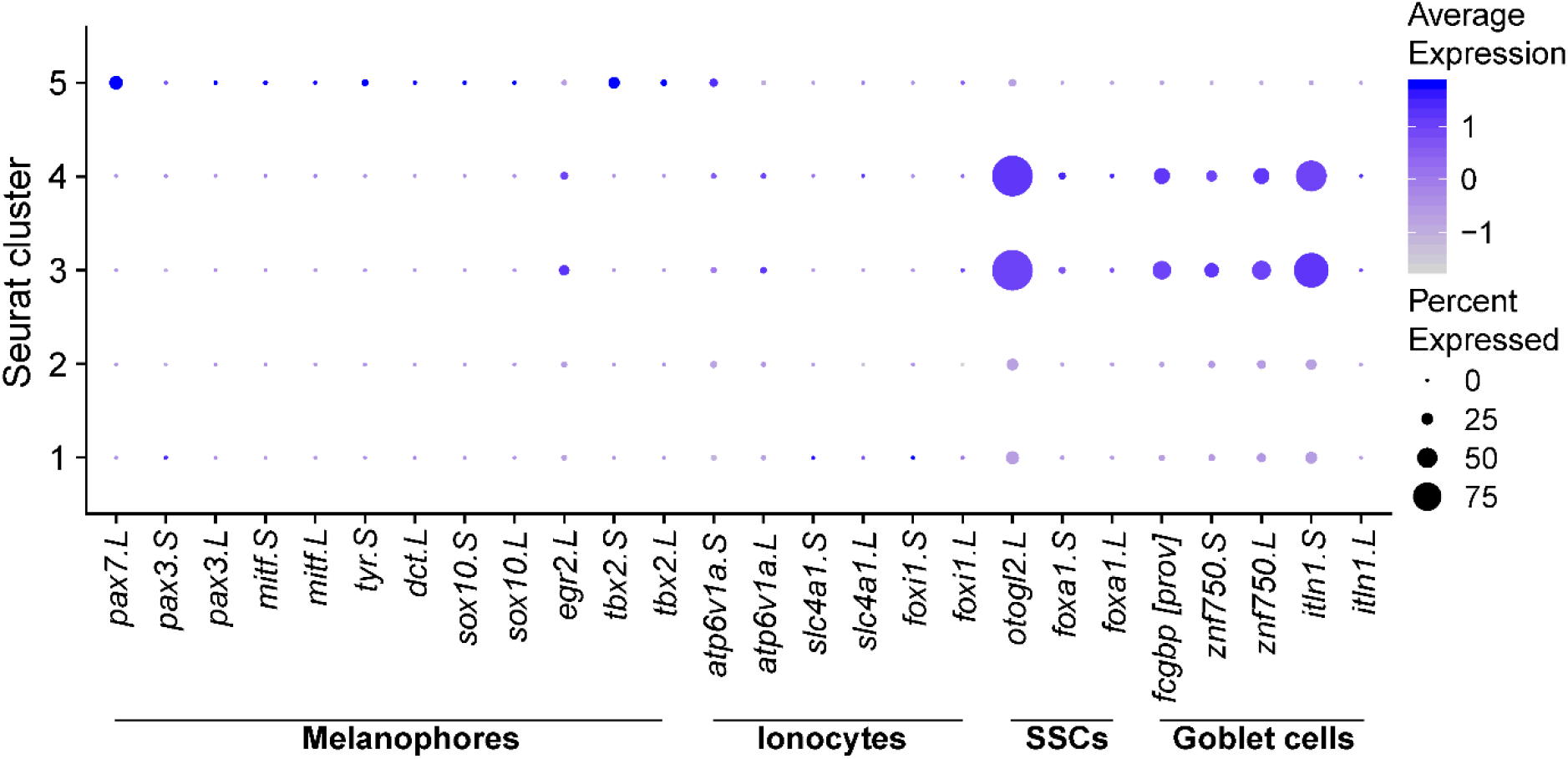
Cells of the apical corneal epithelium display a *Xenopus* skin-like characteristic. Dot plot depicting expression of *Xenopus* skin cell-types (Melanophores; Ionocytes; Small Secretory Cells, SSCs and Goblet cells) marker genes across each cluster in the dataset. Marker genes are listed along the X-axis. Dot diameter depicts the percentage of cluster cells expressing that marker and the color intensity encodes the average expression of that gene among all cells within that cluster, according to the keys.

**Fig. S5.**
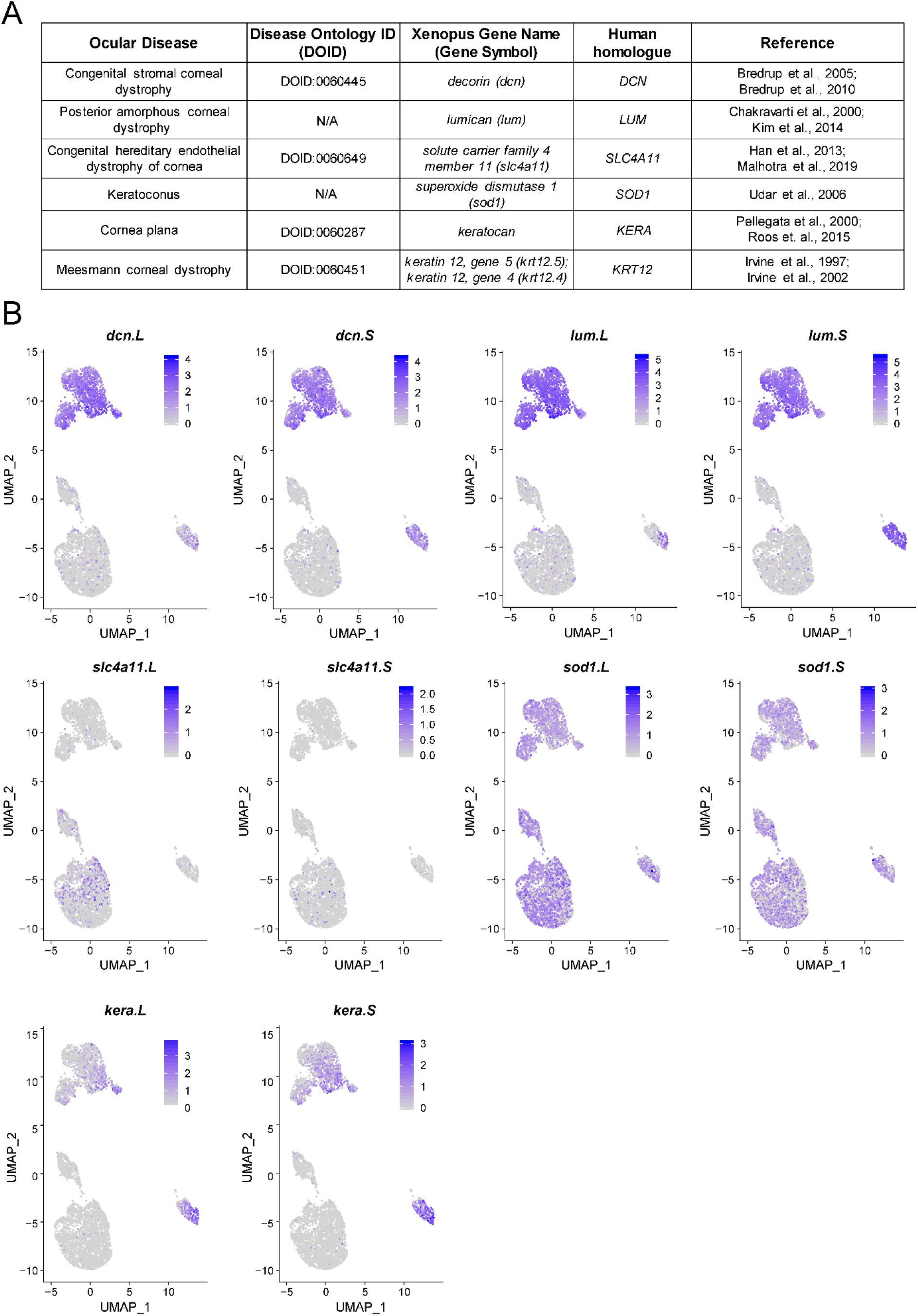
Identification of human corneal dystrophy gene homologues in *Xenopus* corneas. (**A**) Table showing ocular disease condition, Disease Ontology ID (DOID) listed in xenbase.org, *Xenopus* gene names, human homologue and disease literature references. (**B**) Feature plot of *Xenopus* homologues of genes associated with corneal dystrophies identified in our scRNA-seq data. Relative expression is indicated in the keys. Feature plot of *krt12* is shown in **Fig. 4D-E**. Abbreviation: N/A: Not available

---

**Table S1**. List of antibodies used in this study.

**Table S2**. Sequencing metrics for scRNA-seq of corneal epithelial cells from Stage 49-51 Xenopus tadpoles.

**Table S3**. Top 50 marker genes for each cell type cluster, related to **Fig. 4**.

**Table S4**. Enriched gene ontology (GO) terms along pseudo-time, related to **Fig. 6**.

